# Situating the oxytocin receptor gene polymorphisms in the context of structural and connectome-level substrates and association with endogenous oxytocin

**DOI:** 10.1101/2023.08.24.554569

**Authors:** Kutlu Kaya, Deniz Önal, Yasemin Kartal, Murat Timur Budak, Erdem Karabulut, Kader Karlı Oğuz, Bilge Pehlivanoğlu

**Affiliations:** Department of Physiology, Faculty of Medicine, Hacettepe University, Ankara, Turkiye; Department of Physiology, Faculty of Medicine, Balıkesir University, Balıkesir, Turkiye; Department of Physiology, Faculty of Medicine, Kırklareli University, Kırklareli, Turkiye; Department of Biostatistics, Faculty of Medicine, Hacettepe University, Ankara, Turkiye; Department of Radiology, Davis Medical Center, University of California, California, USA

**Keywords:** Oxytocin receptor gene polymorphism, Sex-specific differences, Structural MRI, Resting-state functional connectivity, Structural connectivity, Endogenous oxytocin

## Abstract

Genetic variants in the oxytocin receptor (OXTR) have been linked to individual differences in social behavior, while aberrant oxytocin regulation is associated with increased susceptibility to neuropsychiatric disorders. Recent magnetic resonance imaging (MRI) studies have demonstrated altered brain morphology and connectivity in response to OXTR single nucleotide polymorphisms (SNP) of GG homozygous compared to targeted allele carriers, such as T or A. However, the sex-specific differences in the structural and connectome-level substrates of OXTR genetic variants and their relationship with endogenous oxytocin remain poorly understood. Therefore, we aimed to decompose structural MRI and functional/structural connectivity to identify sex-specific differences among young adults through OXTR SNPs (rs53576, rs1042778, and rs2254298). High-resolution 3D T1-weighted, resting-state functional, and diffusion tensor images were acquired by sixty-one participants who provided blood samples for quantification of endogenous oxytocin concentrations and use for genotyping, followed by grouping with respect to homozygous and targeted allele carriers. We found that men had greater cortical surface area and sub(cortical) gray matter volume in different homozygous and targeted allele carriers. Resting-state functional and structural connectivity (rsFC and SC, respectively) were allocated differently, primarily in temporal and subcortical brain regions. There were also significant sex-specific differences in mean correlations between endogenous oxytocin and SC, whereas rsFC delineated more significant correlations on the node level. Our results provide valuable insights into sex-specific differences in the structural and connectome-level substrates of OXTR SNPs, contributing to understanding the role of oxytocin in socio-emotional processing and highlighting sex-specific differences in genetic and neural mechanisms underlying social behavior.

## Introduction

Oxytocin (OXT) is an evolutionarily well-preserved neuropeptide mainly produced at magnocellular neurons in supraoptic (SON) and paraventricular nuclei (PVN) of the hypothalamus, then transported into the posterior pituitary. OXT is stored in secretory vesicles until released into the peripheral circulation, where OXT encompasses well-known endocrine functions, such as parturition and lactation. In addition, smaller parvocellular neurons in the PVN produce OXT and they all project directly into various brain regions where OXT acts as a neurotransmitter [Ludwig and Leng, 2006]. OXT plays a substantial role in modulating stress responses and social behavior in the mammalian brain [Feldman et al., 2016]. Physiologically, oxytocin poses cell viability, synaptic and structural plasticity in neurons by binding its receptor (i.e., OXTR), which initiates a cascade of intracellular signaling and transcriptional events [Jurek and Neumann, 2018; Pekarek et al., 2020]. The human OXTR gene is located on chromosome 3p25.3, spanning approximately 17 kilobytes, and contains four exons and three introns [Maud et al., 2018]. The expression of OXTR gene is widespread throughout the brain, markedly in olfactory bulbs in addition to the sensory and prefrontal cortex, nucleus accumbens (NAcc), caudate (CAU), putamen (PU), pallidum (PA), hippocampus (HI), amygdala (AMY), hypothalamus, ventral tegmentum, temporal lobe, and anterior cingulate cortex [Bethlehem et al., 2013; Grinevich and Stoop, 2018; Quintana et al., 2019; Yoshida et al., 2009]. Furthermore, OXTR designates hippocampal and parahippocampal gyrus (PaHG) engagement by elevating the expression of neurotrophic factors such as brain-derived neurotrophic factors [Bakos et al., 2018; Dölen et al., 2013; Zhang et al., 2020].

OXT centrally delineates neuroadaptation, including learning and memory, bonding behavior such as maternal and social bonding, and addiction [Carson et al., 2013; Chini et al., 2014; Meyer-Lindenberg et al., 2011; Yamasue et al., 2012]. More specifically, OXT is known for mediating memory for facial recognition, trust, positive emotions (e.g., love and empathy), and prosocial behaviors (e.g., attachment and affiliation) [Kosfeld et al., 2005; McQuaid et al., 2014; Savaskan et al., 2008; Zik and Roberts, 2015]. Therefore, aberrant oxytocin regulation may confer various psychopathologies, including autism spectrum disorder, schizophrenia, and anxiety disorders [Chagnon et al., 2015; Feldman et al., 2016; Meyer-Lindenberg et al., 2011]. Magnetic resonance imaging (MRI) studies have highlighted changes in brain structure and function with respect to genetic variants in OXTR [Baumgartner et al., 2008; Kirsch et al., 2005; Meyer-Lindenberg, 2008]. Single nucleotide polymorphisms (SNPs) in the OXTR gene have been shown to influence gene expression and, thus, socio-emotional behaviors [Feldman et al., 2016]. The altered OXTR genotypes correlate well with changes in local network metrics and functional connectivity between the HI, AMY, medial prefrontal cortex, dorsal anterior cingulate cortex, basal ganglia, and thalamus (TH) [Luo et al., 2020]. Three OXTR SNPs of rs53576, rs1042778, and rs2254298 have been widely investigated to elucidate their influence on human behavior. Consequently, the rs2254298 has been linked to autism spectrum disorder, depression among women, and smaller bilateral AMY volumes [Apter-Levy et al., 2013; Furman et al., 2011; Jacob et al., 2007]. While rs1042778 implies reduced prosocial behaviors, rs53576 indicates reduced empathy and prosocial behaviors, demonstrating that these SNPs have a substantial role in human socio-emotional functioning [Bakermans-Kranenburg and IJzendoorn, 2008; Feldman et al., 2012; Rodrigues et al., 2009]. However, this raises the question of whether these SNPs directly impact functional/structural connectivity or whether it varies with respect to the endogenous OXT concentrations.

Notably, plasma OXT concentrations may demonstrate sex-specific differences in brain functionality since they are robustly associated with central OXT concentrations [Carson et al., 2015]. Because endogenous OXT concentrations in plasma may vary with age, sex and fluctuate in women with respect to the period of the menstrual cycle, plasma OXT levels may lead to ambiguous results on brain structure and function [Plasencia et al., 2019]. Nevertheless, as a surrogate measure of central OXT activity, higher endogenous OXT concentrations were found in response to stressful events in women but not men [Tops et al., 2013; Weisman et al., 2013]. Conversely, social cognitive ability was correlated with plasma OXT concentrations in men [Deuse et al., 2018; Strauss et al., 2019]. Despite the accumulating evidence, the relationship in endogenous OXT concentrations with structural and connectome-level substrates has yet to be disentangled.

The sex-specific differences in the same SNP may provide critical information to comprehend socio-emotional behaviors. Therefore, in this study, we aimed to decompose the influence of OXTR SNPs on structural and connectome-level substrates among young adults. Moreover, we provide comprehensive substrates of the sex-specific relationship in endogenous OXT concentrations and its association with structural/functional connectivity on the node level.

Detailed organization, differential contribution of cortical thickness

Cortical surface area, cortical thickness, and subcortical volumes are measures of brain structure and morphology. They provide valuable information about the organization and development of the brain, and they are often used in neuroimaging studies to study brain differences, development, aging, and various neurological and psychiatric disorders. Cortical surface area, cortical thickness, and subcortical volumes are measures of various aspects of brain structure and morphology that provide insights into brain development, aging, and neurological conditions.

## Methods

### Participants

Sixty-three volunteers enrolled for this study between April 2019 and March 2020 through an advertisement on the university campus. Sixty-one participants (age 21 ± 2, range 18–27 years, 27 women) were included for data analysis. One participant was excluded due to excessive head movement in the functional data, and one was excluded due to artifacts in the diffusion data. Upon explaining the study to the participants who met the inclusion criteria (e.g., no neurological or psychiatric disorders and no contradictions to MRI), they were given sufficient time to decide whether to participate and sign the informed consent. Each participant provided blood samples to measure endogenous OXT concentrations from plasma and to use for genotyping, followed by MRI acquisitions. In addition, we tracked the menstrual cycles of women participants, ensuring that they were in the same phase (days 18–21, luteal phase) for MRI data acquisition and blood testing.

### Quantification of endogenous oxytocin concentrations

The blood samples were centrifuged at 4°C at 1.200 g for 10 min to recover plasma. Upon plasma isolation, samples were aliquoted in several tubes to avoid freeze/thaw cycles, which might degrade the OXT, and stored at –80°C until the biochemical analysis. All samples were assayed within three months of collection to limit OXT degradation during storage. Endogenous OXT concentrations were measured at 450 nm with the SpectraMax plate reader (Molecular Devices, San Jose, CA, USA) from plasma using the ELISA kit (catalog number: E1046Hu, Bioassay Technology Laboratory) with respect to the manufacturer’s instructions. According to the manufacturer, the kit’s sensitivity was 1.06 pg/ml, and the detection range was 2–600 pg/ml.

### Genotyping

The SNPs of rs53576, rs1042778, and rs2254298 from the OXTR gene were selected for genotyping. First, the genomic DNA was extracted by the manufacturer’s instructions using the PureLink Genomic DNA Mini Kit (catalog number: K1820–02, Invitrogen, Carlsbad, CA, USA). The quality of DNA was evaluated by measuring absorbance readings for all samples via a NanoDrop spectrophotometer (Thermo Fisher Scientific, Waltham, MA, USA) and assessed by the A260/280 ratio. The genotyping was then performed for rs53576 (ID: C_3290335_10), rs1042778 (ID: C_7622140_30), and rs2254298 (ID: C_15981334_10) with a commercial TaqMan Genotyping Assay (Applied Biosystems, Foster City, CA, USA). Briefly, the genotyping mix was meticulously prepared in a volume of 10 µl that contained 10 ng of genomic DNA, 0.25 µl of 40× TaqMan Assay mix, and 5 µl of 2× TaqMan Master mix. Upon plate preparation, the genes were genotyped by ABI 7500 Fast Real-Time PCR system (Applied Biosystems, Foster City, CA, USA). Table 1 indicates the demographics and endogenous OXT concentrations of participants that were divided into groups in regard to GG homozygous (SNP^(+)^) and targeted allele carriers (e.g., T or A; SNP^(–)^).

**Table 1.** Demographics and endogenous OXT concentrations of participants stratified by genotype.

| OXTR SNP | Genotype | *n*<br>(M/W) | Age<br>(years) | Endogenous OXT<br>(pg/ml) | *p*<br>(M vs W) |
| --- | --- | --- | --- | --- | --- |
| rs53576<sup>(+)</sup> | GG | 15/14 | 20 ± 2 | 410 ± 297 | 0.83 |
| rs53576<sup>(-)</sup> | GA/AA | 19/13 | 21 ± 2 | 303 ± 219 | 0.64 |
| rs1042778<sup>(+)</sup> | GG | 20/18 | 21 ± 2 | 369 ± 298 | 0.97 |
| rs1042778<sup>(-)</sup> | GT/TT | 14/9 | 21 ± 2 | 346 ± 245 | 0.73 |
| rs2254298<sup>(+)</sup> | GG | 24/18 | 21 ± 2 | 350 ± 284 | 0.28 |
| rs2254298<sup>(-)</sup> | GA/AA | 10/9 | 21 ± 2 | 364 ± 219 | 0.41 |

OXT, oxytocin; OXTR, oxytocin receptor; SNP, single nucleotide polymorphism; M, men; W, women; SNP<sup>(+)</sup>, homozygous; SNP<sup>(-)</sup>, targeted allele carrier; pg, picogram; ml, milliliter; data are presented as mean ± standard deviation.

### MRI data acquisitions

Each participant underwent four neuroimaging scans using a 3-Tesla MRI Scanner equipped with a 32-channel quadrature head coil (Magnetom Trio, Siemens, Erlangen, Germany). The imaging protocol included structural MRI, resting-state functional MRI (rs-fMRI), and diffusion tensor imaging (DTI). An experienced radiologist reviewed structural images on site to confirm the eligibility of the participants during the MRI session.

A high-resolution 3D T1-weighted (W) magnetization-prepared rapid gradient echo (MPRAGE) anatomical images were acquired with the following parameters: repetition time (TR) = 2600 ms, echo time (TE) = 3.02 ms, matrix = 256 × 256, slices = 256, voxel size = 1.0 × 1.0 × 1.0 mm^3^, no intersection gap, slice thickness = 1.00 mm, flip angle (FA) = 8°, field of view (FOV) = 256 mm, and scan time = 7.18 min. Axial T2-W turbo spin-echo (TSE) images were also acquired from each participant for radiologic inspection (TR/TE = 4000/102 ms, slices = 25, voxel size = 0.7 × 0.7 × 5.0 mm^3^, no intersection gap, matrix = 320 × 320, FOV = 210 mm, FA = 130°, scan time = 2.34 min).

As for rs-fMRI, T2*-W images were acquired in a single session with a gradient-echo planar sequence using blood-oxygen-level-dependent (BOLD) contrast for 5 min 48 sec (TR/TE = 2000/35 ms, voxel size = 3.0 × 3.0 × 3.0 mm^3^, without intersection gap, FOV = 192 mm, matrix = 64 × 64, slices = 28; FA = 75°, slice thickness = 3.0 mm) during which participants were asked to remain still and keep their eyes open during the scanning.

DTI scans were acquired using a single shell with 40 independent gradient-encoded directions (*b* = 1000 s/mm^2^) and 10 non-diffusion-weighted (*b* = 0 s/mm^2^) in the axial plane (TR/TE = 10740/102 ms, voxel size = 2.0 × 2.0 × 2.0 mm, FoV = 256 mm, matrix = 128 × 128, slices = 64, without intersection gap, slice thickness = 2.00 mm, scan time = 9.09 min).

### Processing of MRI data

The preprocessing of the 3D T1-W images was performed with the FreeSurfer v7.3.2 (https://surfer.nmr.mgh.harvard.edu) via the autorecon processing, entailing motion correction, intensity normalization, Talairach registration, skull stripping, tessellation of the gray matter/white matter (GM/WM) boundary, segmentation of subcortical WM, automated topology correction, and surface deformation [Fischl et al., 1999; Fischl et al., 2002; Fischl et al., 2004; Fischl and Dale, 2000]. While running the autorecon processing, we implemented an optional step in which several different smoothing kernels from 0 to 25 mm with 5 mm increments of full-width-at-half-maximum (FWHM) were generated to allow reliable group-level analysis. This was followed by concatenating these images into a single image, then spatially smoothing the data with a 10 mm FWHM Gaussian kernel. We visually inspected the reconstruction of each participant and manually corrected through vertex edits or control points in case of major topological inaccuracies. The cortex was parcellated into 34 gyral-based region-of-interest (ROI) per hemisphere with respect to the Desikan-Killiany atlas [Desikan et al., 2006]. We then quantified the total cortical surface area (cSA) of the pial, cortical thickness (CTh), and cortical and subcortical GM volumes.

To preprocess the rs-fMRI data, we used CONN toolbox v21.a (https://www.conn-toolbox.org) and SPM12 (https://www.fil.ion.ucl.ac.uk/spm/) with the following steps to perform the volume-based analysis: removal of 5 initial functional scans to allow for T1-equilibration, realigning and unwarping of functional data using a least squares approach and rigid body transformation, resampling with b-spline interpolation to correct for motion and magnetic susceptibility interactions, slice-timing correction of functional data for temporal misalignment between slices, outlier detection in functional volumes, segmentation into GM, WM, and cerebrospinal fluid (CSF) tissue classes, normalization of the functional and anatomical data into standard Montreal Neurological Institute (MNI) space, and resampling to 2 mm isotropic voxels with a direct normalization procedure. Outlier detection was defined in parameters of the global BOLD signal changes above 5 standard deviations or subject-motion threshold of 0.9 mm scan-to-scan frame-wise displacement. The functional images were spatially smoothed with an 8 mm FWHM Gaussian kernel [Whitfield-Gabrieli and Nieto-Castanon, 2012].

DTI data were preprocessed with the following steps: (i) denoising and removing Gibbs’ ringing artifacts by MRtrix v3.0.4 (https://www.mrtrix.org), (ii) alignment, distortion, and eddy-currents corrections by FSL v6.0.5 (http://www.fmrib.ox.ac.uk/fsl), and (iii) bias field corrections by ANTs v2.4.0 (http://stnava.github.io/ANTs) [Jenkinson et al., 2012; Tournier et al., 2019]. We then created a basis function for each participant to determine the diffusion orientation within each voxel. Since we had single-shell data, we estimated the response function via the *tournier* algorithm for the WM response [Tournier et al., 2013]. Then, fiber orientation distributions (FODs) were estimated to deconvolve the amount of diffusion in each orthogonal direction with the constrained spherical deconvolution method [Behrens et al., 2003; Tournier et al., 2007]. Lastly, we normalized the FODs to allow reliable group-level comparisons.

### Functional connectome construction

Artifact Detection Tools (https://www.nitrc.org/projects/artifact_detect) was used to eliminate the outlier data points through the aforementioned parameters upon BOLD signal preprocessing. A band-pass filter of 0.01–0.1 Hz was applied to reduce interference of physiological noise with linear detrending. The denoising of the functional data included the elimination of WM and CSF noise using aCompCor and the regression of confounding effects such as nuisance regressors, motion correction, and scrubbing [Behzadi et al., 2007]. Upon denoising, voxel-wise BOLD time series were averaged into ROI time series regarding Harvard-Oxford cortical and subcortical atlases. This was followed by the computation of the ROI-to-ROI connectomes through a bivariate analysis of Pearson’s correlation coefficients, resulting in a 105×105 Fisher *z*-transformed adjacency matrix.

### Structural connectome construction

We segmented the anatomical images to employ anatomically-constrained tractography (ACT). We then coregistered anatomical and diffusion images; thereby, both boundaries were robustly aligned. We generated a mask image concerning seed streamlines on the GM/WM interface. ACT is performed through the probabilistic iFOD2 algorithm with the following parameters: ten million streamlines, cut off = 0.06, maximum length = 250 mm, unidirectional tracking, maximum angle in degrees between successive steps = 45 with backtrack option [Tournier et al., 2010]. Additionally, tractograms were optimized for cross-sectional multipliers to counter-balance the whole brain tractography across fixel-based fiber densities and then were filtered to one million streamlines. To reduce the spurious connections, we used average fractional anisotropy between two regions to generate a structural network [Besson et al., 2014]. Lastly, a weighted adjacency matrix (82×82) was generated for each participant (without cerebellum) using the participant’s anatomy obtained by FreeSurfer output and streamlines between each parcellation pair.

### Statistical analyses

The demographic and biochemical characteristics of the participants on a sex basis and for SNP^(+)^ and SNP^(–)^ were compared by the Mann-Whitney U test. We tested sex-specific differences for each SNP in cSA, CTh, and cortical GM volume by performing the non-parametric permutation test with 5.000 iterations and a vertex-wise threshold of 3.0 (*p* < 0.001) for multiple comparisons to avoid inflated false positives [Greve and Fischl, 2018]. The significant clusters were then evaluated with cluster-wise *p*-values (*CWP*) < 0.05. To compare differences in subcortical GM volumes, we performed ROI analysis using the non-parametric Mann-Whitney U test with the Bonferroni correction for multiple comparisons.

The individual ROIs were tested for resting-state functional connectivity (rsFC) with the false discovery rate (FDR) correction for multiple comparisons. The structural connectivity (SC) was tested by performing the network-based statistics with a *t*-threshold of 3.5 and 5.000 permutations with the family-wise error (FWE) correction [Zalesky et al., 2010].

To explore the relationship between endogenous OXT concentrations and network-level connectivity, Pearson’s correlation analysis was performed at each node using the arithmetic mean method. To reveal sex-specific total node correlation differences, we performed the independent *t*-test, where statistical significance was evaluated employing a two-tail *t*-distribution with a 95% confidence interval. The data are presented as mean with standard deviation (SD), and *p*-values < 0.05 were considered statistically significant, including post-correction with the corresponding test (*p*_corrected_ = Bonferroni, FDR, or FWE).

## Results

### Cortical surface area, (sub)cortical GM volumes, and thickness

Figure 1 indicates sex-specific cSA differences stratified by OXTR SNPs. The vertex-wise analysis for the rs53576^(+)^ showed that men had larger cSA in the left middle temporal (MTG, *CWP* = 0.04) and fusiform gyri (FG, *CWP* = 0.04). We also found larger cSA in the right inferior temporal gyrus (ITG, *CWP* = 0.03) for the rs1042778^(+)^ men. For the rs2254298^(+)^ men, we found larger cSA in the left MTG (*CWP* < 0.001), precentral (PreCG, *CWP* = 0.002), and superior parietal gyri (SPG, *CWP* = 0.01). Furthermore, there was a larger cSA in the right ITG (*CWP* < 0.001), superior frontal gyrus (SFG, *CWP* = 0.02), and PreCG (*CWP* = 0.04) for the rs2254298^(+)^ men. The sex-specific differences in cSA regarding the targeted allele carriers were also observed in all SNPs. The rs53576^(–)^ men had larger cSA in the left lateral occipital cortex (LOC, *CWP* = 0.01), left ITG (*CWP* = 0.04), bilateral PreCG (left: *CWP* < 0.001 and right: *CWP* = 0.03), and bilateral SPG (*CWP* = 0.01 for both). The rs1042778^(–)^ men had larger cSA in the left PreCG (*CWP* = 0.01), MTG (*CWP* = 0.04), LOC (*CWP* < 0.001), and the right cuneus (CU, *CWP* = 0.01). We found larger cSA in the left FG (*CWP* = 0.01) and the right CU (*CWP* = 0.005) for the rs2254298^(–)^ men (see Supplementary Table S1 for details).

**Figure 1.**
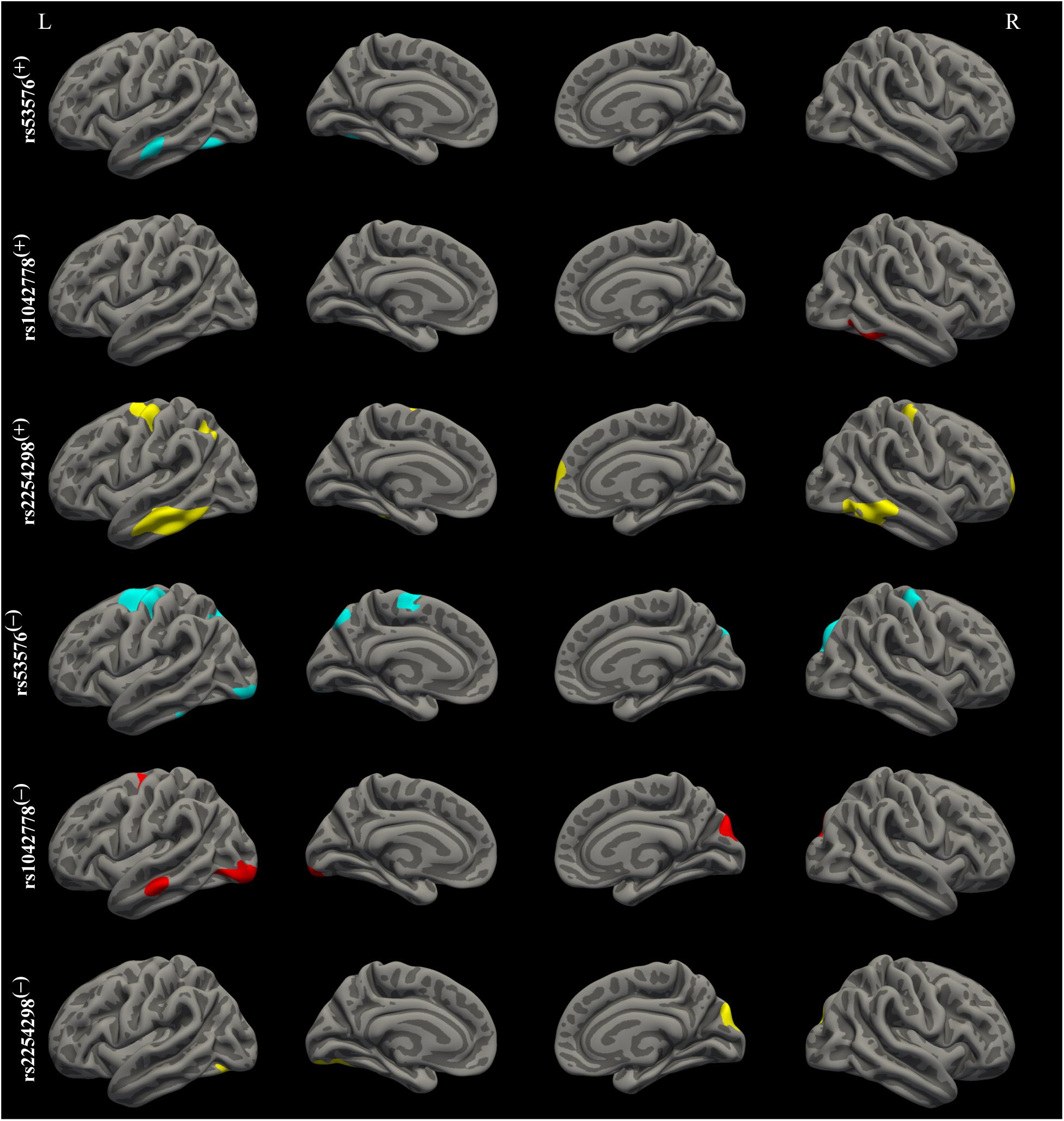
Sex-specific cortical surface area (cSA) differences stratified by single nucleotide polymorphisms (SNPs). Compared to women, the cSA for homozygous (SNP^(+)^) and targeted allele carriers (SNP^(–)^) in men. The SNP rs53576 is in turquoise, the SNP rs1042778 is in red, and the SNP rs2254298 is in yellow. The significant clusters were assessed with cluster-wise *p*-values < 0.05. Lateral and medial views are demonstrated in the left (L) and right (R) hemispheres.

As indicated in Figure 2, there were also sex-specific differences in cortical and subcortical GM volumes for the SNPs. For the cortical regions, the volume of the left ITG had significantly greater in men for the rs53576^(+)^ (*CWP* = 0.01) and rs2254298^(+)^ (*CWP* = 0.007). Furthermore, men had greater volumes of the right SFG (*CWP* = 0.02), LOC (*CWP* = 0.02), and FG (*CWP* = 0.04) for the rs2254298^(+)^. However, there were no significant differences in the cortical volume for the rs1042778^(+)^. For the targeted allele carriers, bilateral PreCG (left: *CWP* = 0.03 and right: *CWP* = 0.001) and left ITG (*CWP* = 0.02) had a greater volume for the rs53576^(–)^ men. In addition, men had significantly greater volumes of the left SFG and the right SPG for the rs53576^(–)^ (*CWP* = 0.04 and *CWP* = 0.001, respectively) and rs1042778^(–)^ (*CWP* = 0.03 and *CWP* = 0.007, respectively). Finally, the volumes of the left FG (*CWP* = 0.002) and bilateral precuneus (PreCU, *CWP* = 0.03 for both) were significantly greater for the rs2254298^(–)^ men (see Supplementary Table S2 for details).

**Figure 2.**
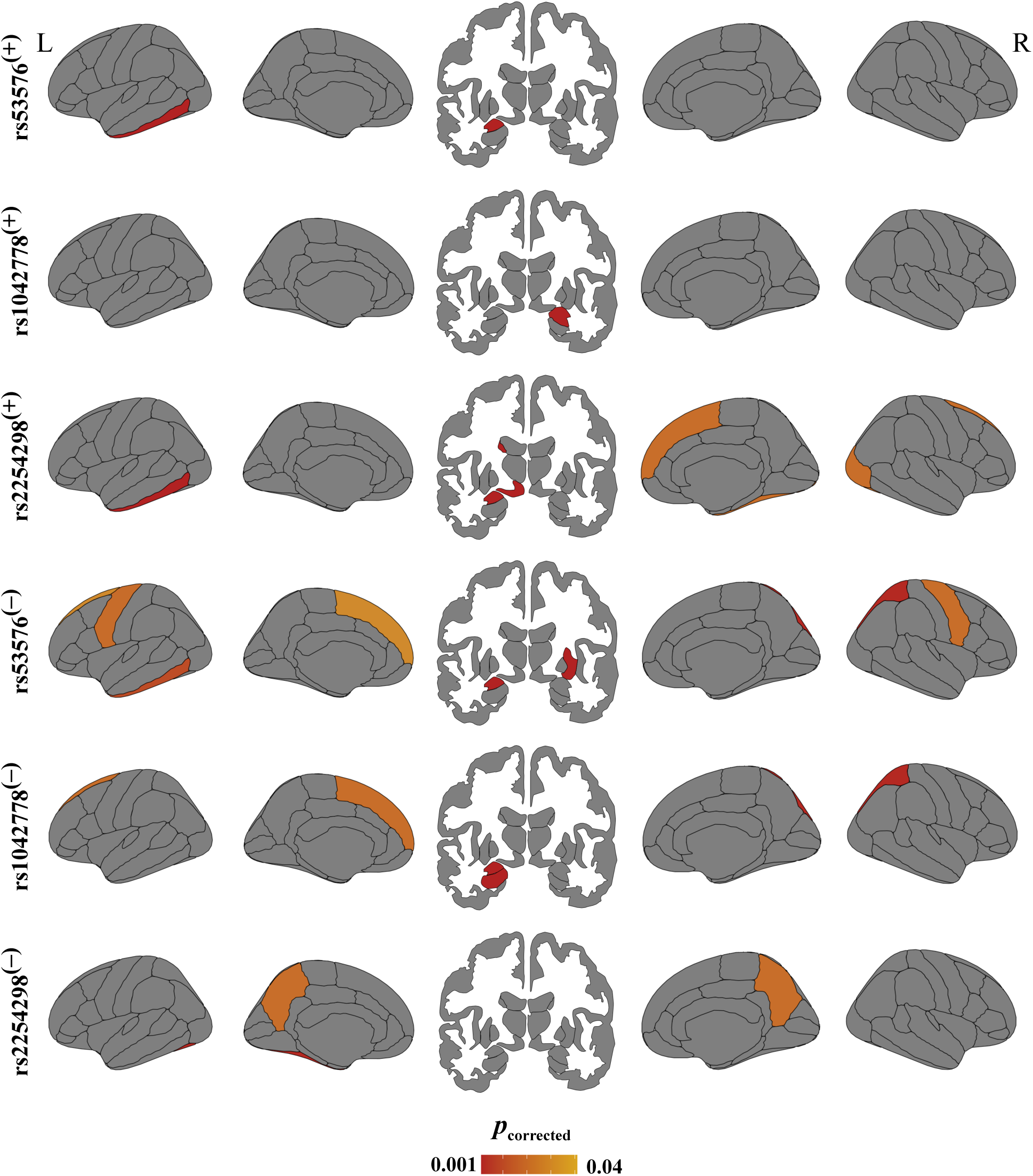
Sex-specific cortical and subcortical gray matter (GM) volume differences stratified by single nucleotide polymorphisms (SNPs). Compared to women, the GM volume for homozygous (SNP^(+)^) and targeted allele carriers (SNP^(–)^) in men. Significant differences were shown after correcting for multiple comparisons (*p*_corrected_). Lateral/medial views and subcortical regions are demonstrated in the left (L) and right (R) hemispheres. Figures were ggseg v1.5.3 (https://github.com/ggseg/ggseg) in *R* v4.2.0.

As for the subcortical volumes in homozygous SNPs, men had a significantly greater volume of the left AMY for the rs53576^(+)^ (*p*_corrected_ = 0.006) and rs2254298^(+)^ (*p*_corrected_ = 0.003). The same difference was in the right AMY for the rs1042778^(+)^ (*p*_corrected_ = 0.001). Furthermore, men had a significantly greater volume of the left caudate (CAU) and ventral diencephalon (vDC) for the rs2254298^(+)^ (*p*_corrected_ = 0.004 and *p*_corrected_ < 0.01, respectively). For the targeted allele carriers, men had a significantly greater volume left the AMY for the rs53576^(–)^ (*p*_corrected_ = 0.005) and rs1042778^(–)^ (*p*_corrected_ = 0.01). Moreover, the volume of the right putamen (PU) was greater for the rs53576^(–)^ men (*p*_corrected_ = 0.003) and the left HI for the rs1042778^(–)^ men (*p*_corrected_ = 0.001). However, there were no significant sex-specific differences in subcortical volumes for rs2254298^(–)^ (see Supplementary Table S3 for details).

### Resting-state functional connectivity

The rsFC analysis revealed significant sex-specific differences with respect to homozygous SNPs. For the rs53576^(+)^, we found that men had a positive correlation between the right CU and the left lingual gyrus (LG, *t* = 4.51, *p*_corrected_ = 0.01), while there was a negative correlation between the left frontal operculum (FO) and the right postcentral gyrus (PostCG, *t* = –4.07, *p*_corrected_ = 0.04, Figure 3a–d). As for the rs1042778^(+)^, men had a negative correlation between the right SFG and the right posterior MTG (*t* = –4.27, *p*_corrected_ = 0.04, Figure 3b–d). Moreover, there were more differences in the rs2254298^(+)^ than in other homozygous SNPs. We found negative correlations between the left temporal pole (TP) and the left middle frontal gyrus (MFG, *t* = –3.96, *p*_corrected_ = 0.04), the left anterior MTG and the right posterior temporal fusiform cortex (TFusC, *t* = –4.05, *p*_corrected_ = 0.02), the left anterior MTG and the right anterior PaHG (*t* = –4.04, *p*_corrected_ = 0.02) for men. Additionally, a positive correlation existed between the left anterior supramarginal gyrus (SMG) and the right AMY (*t* = 3.96, *p*_corrected_ = 0.04, Figure 3c–d) for men.

**Figure 3.**
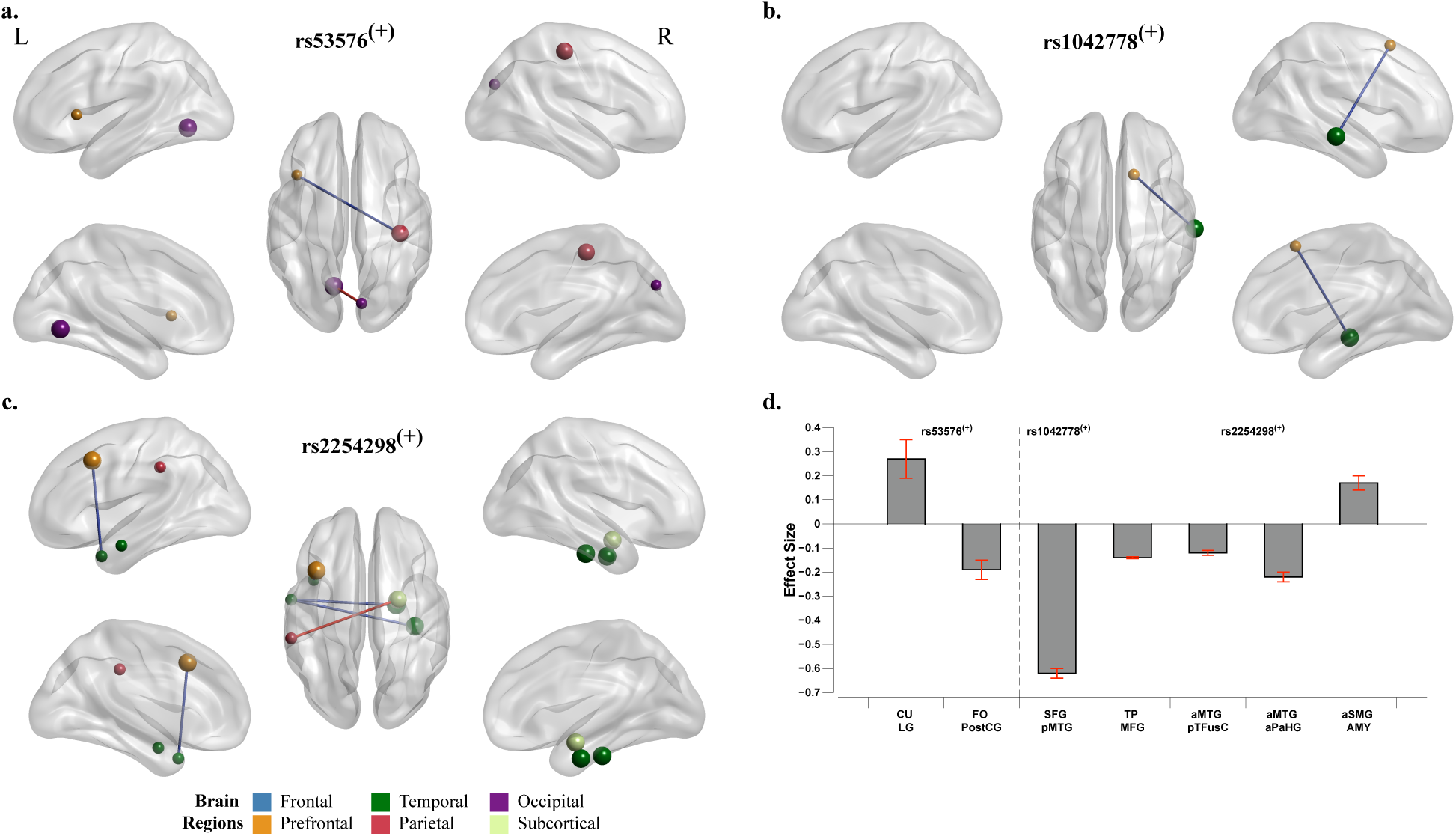
Sex-specific resting-state functional connectivity (rsFC) differences stratified by homozygous single nucleotide polymorphisms (SNP^(+)^). The rsFC for rs53576^(+)^ **(a)**, rs1042778^(+)^ **(b)**, and rs2254298^(+)^ **(c)** in men compared to women. The blue and red lines indicate negative and positive correlations, respectively. The size of nodes represents *t*-scores and colors their location in the brain region both in the left (L) and right (R) hemispheres. Bar graphs indicate effect sizes (mean connectivity scores) at significant pairs **(d)**. Significant correlations were illustrated after false-discovery rate correction. CU, cuneus; LG, lingual gyrus; FO, frontal operculum; PostCG, postcentral gyrus; SFG, superior frontal gyrus; pMTG, posterior middle temporal gyrus; TP, temporal pole; MFG, middle frontal gyrus; aMTG, anterior middle temporal gyrus; pTFusC, posterior temporal fusiform gyrus; aPaHG, anterior parahippocampal cortex; aSMG, anterior supramarginal gyrus; AMY, amygdala. Figures **(a–c)** were created using BrainNet Viewer (https://www.nitrc.org/projects/bnv/).

For the targeted allele carriers, we found negative correlations between the left TP and the left MFG (*t* = –4.13, *p*_corrected_ = 0.03), the right temporal occipital fusiform cortex (TOFusC) and the left Heschl’s gyrus (HG, *t* = –4.10, *p*_corrected_ = 0.04) for the rs53576^(–)^ men. In addition, a positive correlation was found between the left anterior PaHG and the left CAU in the rs53576^(–)^ men (*t* = 4.32, *p*_corrected_ = 0.02, Figure 4a). As for the rs1042778^(–)^ men, negative correlations were found between the right insula (IN) and the left posterior ITG (*t* = –4.16, *p*_corrected_ = 0.02), the left anterior PaHG and the right HI (*t* = –4.15, *p*_corrected_ = 0.03), the left anterior TFusC and the right TH (*t* = – 3.98, *p*_corrected_ = 0.04). Furthermore, there were also positive correlations between the right inferior frontal gyrus pars opercularis (IFG.oper) and the right MFG (*t* = 3.86, *p*_corrected_ = 0.03), the left anterior SMG and the right AMY (*t* = 4.57, *p*_corrected_ = 0.01) in the rs1042778^(–)^ men (Figure 4b). However, there were no significant sex-specific differences for the rs2254298^(–)^.

**Figure 4.**
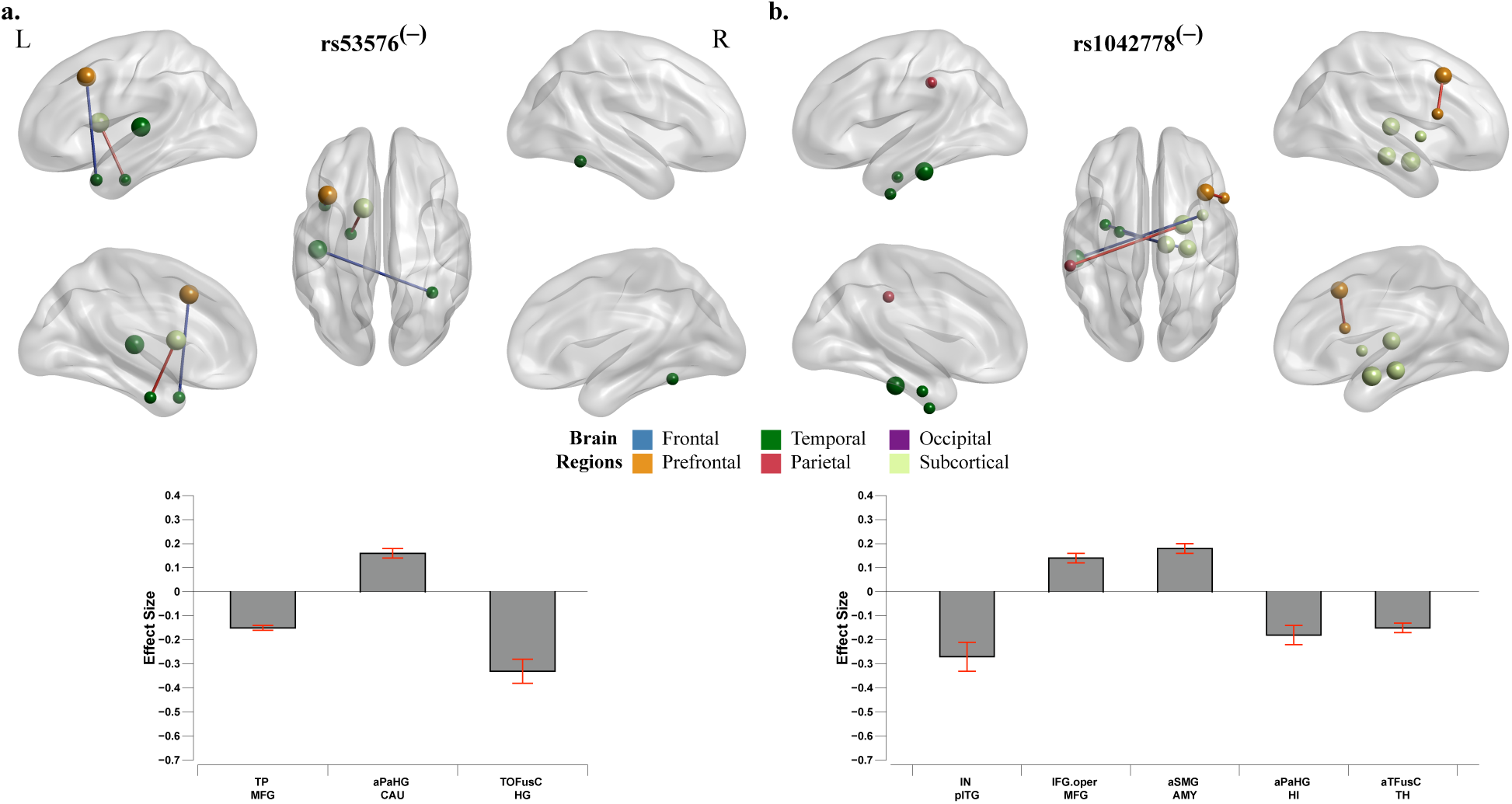
Sex-specific resting-state functional connectivity (rsFC) differences stratified by targeted allele carrier single nucleotide polymorphisms (SNP^(–)^). The rsFC for rs53576^(–)^ **(a)** and rs1042778^(–)^ **(b)** in men compared to women. The blue and red lines indicate negative and positive correlations, respectively. The size of nodes represents *t*-scores and colors their location in the brain region both in the left (L) and right (R) hemispheres. Bar graphs indicate effect sizes (mean connectivity scores) at significant pairs. Significant correlations were illustrated after false-discovery rate correction. TP, temporal pole; MFG, middle frontal gyrus; aPaHG, anterior parahippocampal gyrus; CAU, caudate; TOFusC, temporal occipital fusiform cortex; HG, Heschl’s gyrus; IN, insula; pITG, posterior inferior temporal gyrus; IFG.oper, inferior frontal gyrus pars opercularis; aSMG, anterior supramarginal gyrus; AMY, amygdala; HI, hippocampus; aTFusC, anterior temporal fusiform cortex; TH, thalamus. Top figures were created using BrainNet Viewer (https://www.nitrc.org/projects/bnv/).

### Structural connectivity

Figure 5 shows the significant sex-specific differences in SC. For rs53576^(+)^ and rs53576^(–)^, fewer SC differences were found. The rs53576^(+)^ women had greater SC between the right HI and left PostCG (*t* = 3.71) and left PreCG (*t* = 3.72, *p*_corrected_ = 0.01). Moreover, the right HG had greater SC with left IN (*t* = 3.81) and left HI (*t* = 4.07, *p*_corrected_ = 0.01) in women (Figure 5a). The rs53576^(–)^ women had greater SC between the left entorhinal cortex (EC) and right pallidum (PA, *t* = 4.35) and right AMY (*t* = 4.90) and also between the left PaHG and right PA (*t* = 4.71, *p*_corrected_ = 0.04, Figure 5c).

**Figure 5.**
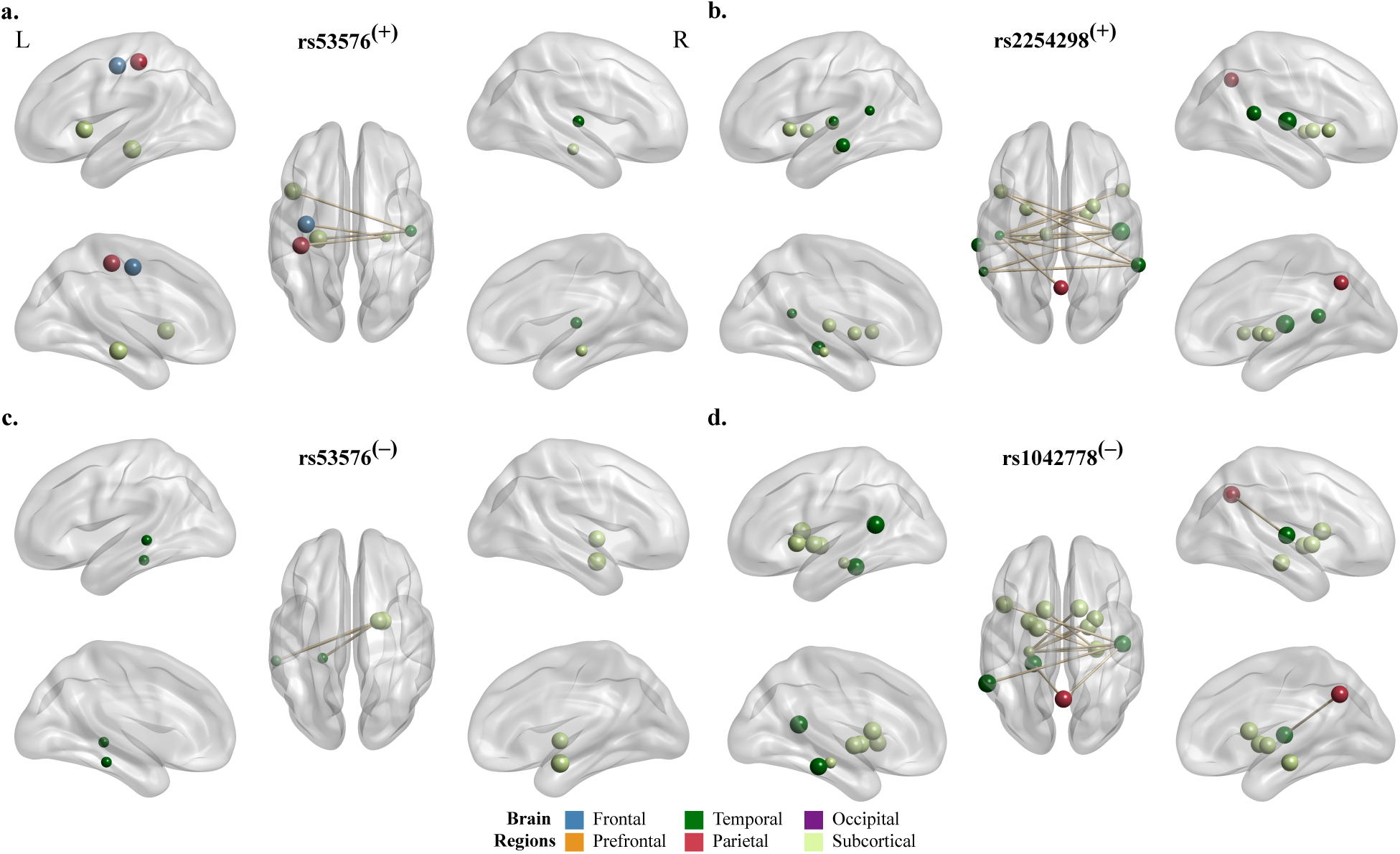
Sex-specific structural connectivity (SC) differences stratified by single nucleotide polymorphisms (SNPs). The SC in homozygous SNPs for rs53576^(+)^ **(a)** and rs2254298^(+)^ **(b)**, and in targeted allele carrier SNPs rs53576^(–)^ **(c)** and rs1042778^(–)^ **(d)** in women compared to men. The significant correlations are indicated with light brown lines after family-wise error correction. The size of nodes represents *t*-scores and colors their location in the brain region both in the left (L) and right (R) hemispheres. Figures were created using BrainNet Viewer (https://www.nitrc.org/projects/bnv/).

The higher SC was observed for rs2254298^(+)^ and rs1042778^(–)^ (Figure 5b and d, respectively), delineating common regions where SC was more allocated. For both SNPs, women had greater SC between the left STG and right HG (rs2254298^(+)^: *t* = 5.66, and rs1042778^(–)^: *t* = 3.95, *p*_corrected_ = 0.002 for both). The left HI was also significantly connected with the right PU (rs2254298^(+)^: *t* = 4.19 and rs1042778^(–)^: *t* = 3.78), right PA (rs2254298^(+)^: *t* = 3.73 and rs1042778^(–)^: *t* = 3.72), and right HG (rs2254298^(+)^: *t* = 4.24 and rs1042778^(–)^: *t* = 3.73, *p*_corrected_ = 0.002 for all) in women. Furthermore, the right PreCU was more structurally connected with the left HG for rs2254298^(+)^ women (*t* = 4.11) while connected with the right HG for rs1042778^(–)^ women (*t* = 3.74, *p*_corrected_ = 0.002 for both). In addition, further SC was observed in rs2254298^(+)^ women than in rs1042778^(–)^ (see Supplementary Table S4 for details).

### The relationship between endogenous OXT concentrations and functional/structural connectivity

Table 2 shows the correlation scores between endogenous OXT concentrations and structural/functional connectivity. There were no significant sex-specific differences in mean correlations between endogenous OXT concentrations and rsFC for any SNPs (Figure 6a). Exploring the association between endogenous OXT concentrations and each connectivity node, there were more significant correlations in rsFC on the node level than that of SC. We found a significant correlation in 15 rsFC nodes for rs1042778^(+)^ and rs2254298^(+)^ in men, primarily in the occipital and temporal brain regions. Furthermore, men had more significant correlations for all targeted allele carriers in 19 nodes (Figure 6b, top). As for women, there were 21 significant rsFC nodes for all homozygous SNPs and 11 significant nodes for all targeted allele carriers (Figure 6b, bottom). The relationship between endogenous OXT concentrations and rsFC was mainly positive in men (19 nodes) and women (27 nodes).

**Figure 6.**
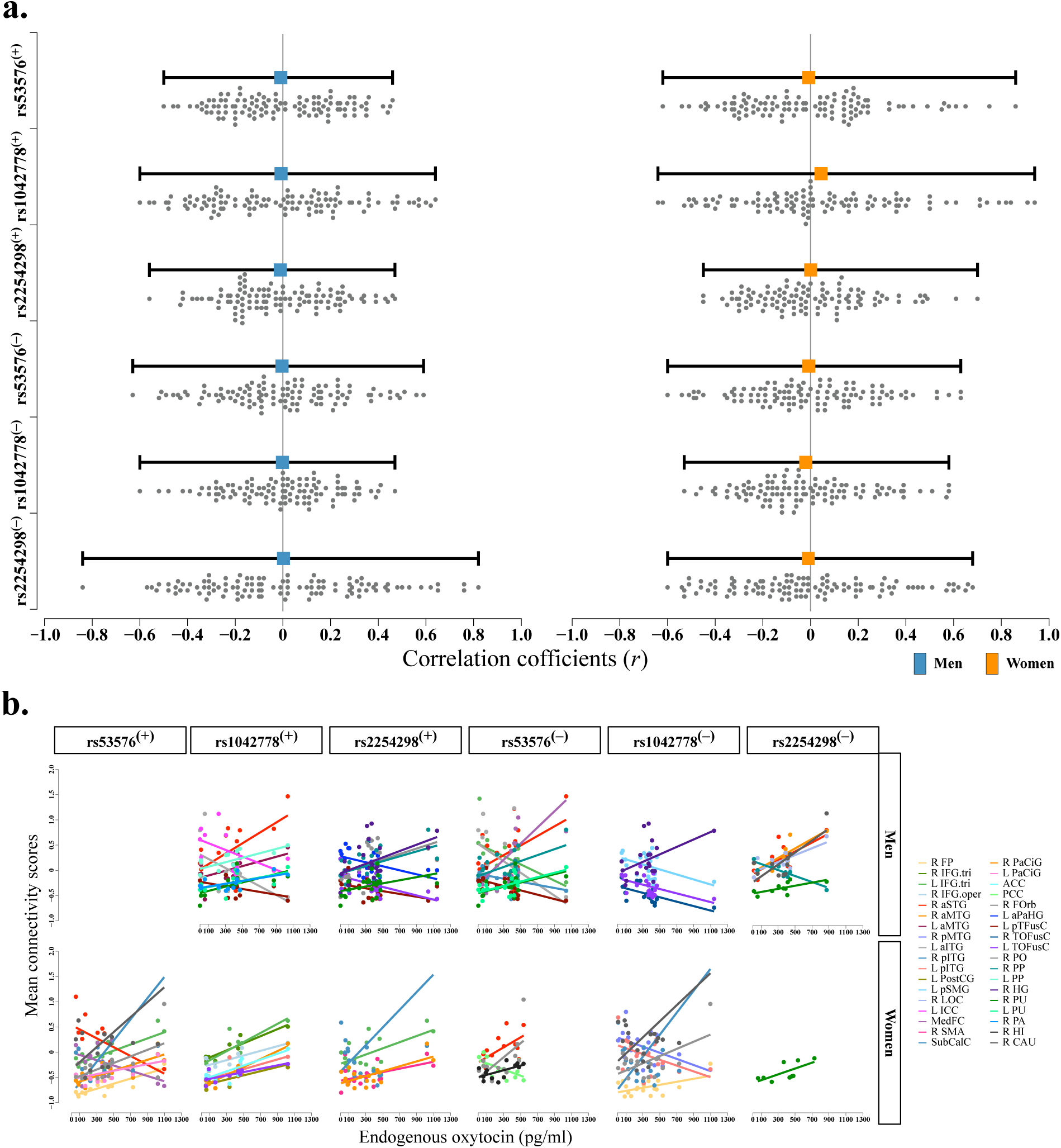
**(a)** Sex-specific differences in the correlation coefficients (*r*) between resting-state functional connectivity (rsFC) and endogenous oxytocin (OXT) concentrations. **(b)** Significant correlations between rsFC and endogenous OXT concentrations on the nodes level for single nucleotide polymorphisms (SNPs) in the homozygous (SNP^(+)^) and targeted allele carriers (SNP^(–)^). Each node is displayed in a different color. L, left; R, right; FP, frontal pole; IFG.tri, inferior frontal gyrus pars triangularis; IFG.oper, inferior frontal gyrus pars opercularis; aSTG, anterior superior temporal gyrus; aMTG, anterior middle temporal gyrus; pMTG, posterior middle temporal gyrus; aITG, anterior inferior temporal gyrus; pITG, posterior inferior temporal gyrus; PostCG, postcentral gyrus; pSMG, posterior supramarginal gyrus; LOC, lateral occipital cortex; ICC, intracalcarine cortex; MedFC, frontal medial cortex; SMA, supplementary motor area; SubCalC, subcallosal cortex; PaCiG, paracingulate gyrus; ACC, anterior cingulate cortex; PCC, posterior cingulate cortex; FOrb, frontal orbital cortex; aPaHG, anterior parahippocampal gyrus; pTFusC, posterior temporal fusiform cortex; TOFusC, temporo-occipital fusiform cortex; PO, parietal operculum; PP, planum polare; HG, Heschl’s gyrus; PU, putamen; PA, pallidum; HI, hippocampus; CAU, caudate.

**Table 2.** Mean correlation scores between endogenous OXT concentrations and functional/structural connectivity in women and men stratified by genotype.

| SNP | Connectivity | Women | Men | df | t | p |
| --- | --- | --- | --- | --- | --- | --- |
| rs53576<sup>(+)</sup> | SC | -0.09 ± 0.20 | -0.02 ± 0.27 | 148 | 2.03 | 0.04 |
|  | rsFC | -0.01 ± 0.29 | -0.01 ± 0.23 | 208 | -0.02 | 0.98 |
| rs1042778<sup>(+)</sup> | SC | -0.17 ± 0.25 | 0.08 ± 0.27 | 162 | 6.10 | < 0.001 |
|  | rsFC | 0.04 ± 0.35 | -0.01 ± 0.31 | 208 | -1.13 | 0.26 |
| rs2254298<sup>(+)</sup> | SC | -0.10 ± 0.19 | 0.04 ± 0.18 | 162 | 4.84 | < 0.001 |
|  | rsFC | 0.001 ± 0.23 | -0.01 ± 0.22 | 208 | -0.34 | 0.73 |
| rs53576<sup>(-)</sup> | SC | -0.05 ± 0.33 | 0.08 ± 0.19 | 128 | 3.09 | 0.002 |
|  | rsFC | -0.01 ± 0.27 | -0.01 ± 0.26 | 208 | 0.11 | 0.91 |
| rs1042778<sup>(-)</sup> | SC | -0.01 ± 0.20 | -0.03 ± 0.20 | 162 | -0.67 | 0.51 |
|  | rsFC | -0.02 ± 0.24 | -0.01 ± 0.22 | 208 | 0.55 | 0.58 |
| rs2254298<sup>(-)</sup> | SC | 0.13 ± 0.31 | -0.02 ± 0.30 | 162 | -3.19 | 0.002 |
|  | rsFC | -0.01 ± 0.31 | 0.01 ± 0.34 | 208 | 0.27 | 0.79 |

OXT, oxytocin; SNP, single nucleotide polymorphism; SNP<sup>(+)</sup>, homozygous allele; SNP<sup>(-)</sup> targeted allele carrier; SC, structural connectivity; rsFC, resting-state functional connectivity; df, degree of freedom; correlation coefficients (*r*) are presented as mean ± standard deviation.

We found significant sex-specific differences between endogenous OXT concentrations and SC in all SNPs except rs1042778^(–)^. For SNPs rs1042778^(+)^, rs2254298^(+)^, and rs53576^(–)^, women had significantly decreased mean correlations than men. This was in contrast to rs2254298^(–)^, where men had significantly decreased mean correlations than women. Although there were negative correlations for the rs53576^(+)^ in both men and women, the mean correlations were significantly lower in women than in men (Figure 7a, *p* < 0.05 for all).

**Figure 7.**
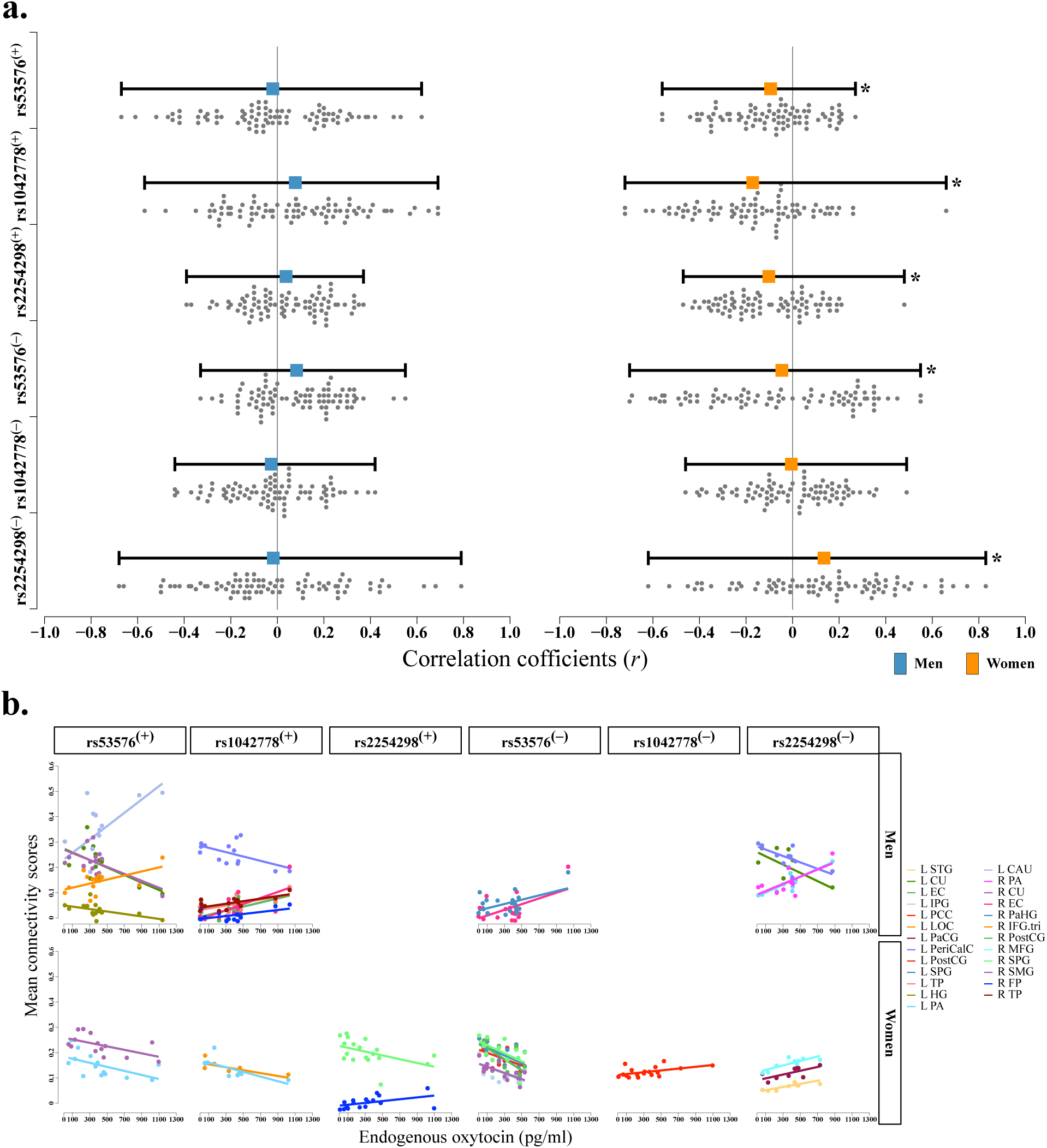
**(a)** Sex-specific differences in the correlation coefficients (*r*) between structural connectivity (SC) and endogenous oxytocin (OXT) concentrations. **(b)** Significant correlations between SC and endogenous OXT concentrations on the nodes level for single nucleotide polymorphisms (SNPs) in the homozygous (SNP^(+)^) and targeted allele carriers (SNP^(–)^). Each node is displayed in a different color. * *p* < 0.05 L, left; R, right; STG, superior temporal gyrus; CU, cuneus; EC, entorhinal cortex; IPG, inferior parietal gyrus; PCC, posterior cingulate cortex; LOC, lateral occipital cortex; PaCG, paracentral gyrus; PeriCalC, pericalcarine cortex; SPG, superior parietal gyrus; TP, temporal pole; HG, Heschl’s gyrus; PA, pallidum; CAU, caudate; PaHG, parahippocampal gyrus; IFG.tri, inferior frontal gyrus pars triangularis; PostCG, postcentral gyrus; MFG, middle frontal gyrus; SMG, supramarginal gyrus; FP, frontal pole.

Delineating the association between endogenous OXT concentrations and SC on the nodes level, we found a significant correlation in 11 nodes for rs53576^(+)^ and rs1042778^(+)^ in men, primarily in the occipital and temporal brain regions as in rsFC. However, there were fewer correlations for rs53576^(–)^ and rs2254298^(–)^ men (6 SC nodes). There was no significant correlation at the SC nodes for rs2254298^(+)^ and rs1042778^(–)^ men (Figure 7b, top).

For women, there was a significant correlation in 6 SC nodes for all homozygous SNPs, primarily located in the parietal, temporal, and subcortical brain regions. Moreover, there was a significant correlation in 10 SC nodes for all targeted allele carriers. Interestingly, all 6 correlations were negative for rs53576^(–)^ women located in bilateral PostCG (left: *r* = –0.56, *p* = 0.04 and right: *r* = –0.69, *p* = 0.009), bilateral SPG (left: *r* = –0.59, *p* = 0.03 and right: *r* = –0.57, *p* = 0.04), left IPG (*r* = –0.61, *p* = 0.03), and right SMG (*r* = –0.70, *p* = 0.008, Figure 6b, bottom). While there were 2 significant positive correlations for rs53576^(–)^ men in the left EC (*r* = 0.55, *p* = 0.01) and the right PaHG (*r* = 0.50, *p* = 0.03, Figure 7b, top).

## Discussion

We utilized surface-based whole brain and functional/structural connectivity analyses to delineate sex-specific differences in young adults through OXTR SNPs. We also explored the sex-specific differences in endogenous OXT concentration and its relation with functional/structural connectivity. The present study demonstrated that different homozygous and targeted allele carriers (1) revealed greater cSA and sub(cortical) GM volume in men, (2) allocated differently in rsFC and SC, primarily located in temporal and subcortical brain regions, and (3) had significant sex-specific differences in mean correlations between endogenous OXT concentrations and SC, whereas rsFC delineated more significant correlations on the node level.

Genetic variants in the OXTR gene can lead to aberrant stress management, altered social behaviors, and deterioration in empathy [Rodrigues et al., 2009; Sicorello et al., 2020]. Moreover, perturbations in oxytocin regulation can yield in psychiatric disorders [Dadds et al., 2014; Mitre et al., 2016]. Indeed, there are subtle morphological and connectome-level changes in the temporal regions and limbic system, causing maladaptive and psychotic traits [Aspé-Sánchez et al., 2016; Israel et al., 2008; Luo et al., 2020]. This straightforward relation is partially determined by neuroanatomy and OXTR gene polymorphism. Previous studies using genetics and functional MRI have shown that OXT influences the structure and function of the human hypothalamic-limbic system [Quintana et al., 2019; Tost et al., 2010]. Specifically, volume-based morphology (VBM) studies have highlighted the influence of OXT on the limbic system, particularly the AMY [Furman et al., 2011]. However, the VBM methodology might produce inflated GM differences through inaccurate segmentation and registration due to the partial volume effect [Callaert et al., 2014; Kennedy et al., 2009]. Therefore, we grasped surface-based analysis given that the results of VBM may not overlap with brain regions in response to cognition [Gerrits et al., 2016].

In this study, we observed sex-specific volume differences in AMY for all SNPs except for rs2254298^(–)^. As the AMY plays a crucial role in emotional processing and the regulation of emotions, the OXTR SNPs may contribute to structural differences in the AMY, potentially influencing emotional responses and social behaviors differently in men compared to women [Tost et al., 2010; Waller et al., 2016]. Although our findings align with previous studies, there is a disparity for rs2254298^(–)^, where previous research reported greater AMY volume in the presence of the A allele among adults [Inoue et al., 2010]. This might arise from methodological differences, as this previous study compared homozygous and targeted allele carriers regardless of sex. Nevertheless, our results imply the potential influence of OXTR polymorphisms on AMY, occupying more volume with respect to OXTR SNPs and hence more functioning in emotional responses and social behavior. Besides, additional subcortical GM volume findings on OXTR SNPs demonstrate how specific genetic variations may contribute to subcortical volume differences, particularly in regions such as the AMY, CAU, vDC, PU, and HI. As these regions are involved in emotion regulation, reward processing, motor control, and memory formation, our results suggest that sex-specific variations in several functions related to the implicated regions [Cahill et al., 2001; Canli et al., 2002; Hamann et al., 2004].

Similarly, homozygous men had larger cSA in temporal regions for all SNPs, signifying greater functioning in language and semantic memory processing, visual perception, and somatomotor functions [Onitsuka et al., 2004]. Interestingly, the rs2254298^(+)^ men showed additional cSA differences involving motor control, spatial orientation, and higher cognitive functions. Moreover, the rs53576^(–)^ and rs1042778^(–)^ men indicated involvement in motor control, visual processing, language, and semantic memory processing, whereas the rs2254298^(–)^ men solely posed involvement in visual processing. The differences in temporal and subcortical regions might demonstrate the combined effect of OXTR polymorphism because subcortical regions, especially AMY, manage information processing between the prefrontal-temporal association cortices and the hypothalamus [Rajmohan and Mohandas, 2007]. Furthermore, the cortical GM volume differences mostly did not accompany the corresponding cSA differences in homozygous and targeted allele carriers even though some SNPs were accompanied (i.e., rs53576^(–)^ and rs2254298^(+)^). Given that the cortical GM volume is ultimately determined by cSA and CTh, the incongruity between cSA and cortical GM volume might be due to the insignificance of CTh [Winkler et al., 2010]. Regarding the association between cSA and CTh, it has been suggested that their cortical development occurs independently of each other and is uncorrelated genetically [Geschwind and Rakic, 2013; Panizzon et al., 2009].

Overall, considering the combined results of cSA and sub(cortical) GM volume differences, several patterns emerge regarding the influence of OXTR SNP and sex-specific differences in cognitive functions and appear to play a role in shaping brain structure and contributing to sex-specific differences. These findings are in line with previous studies indicating higher cSA and cortical volumes in men and higher WM tract complexity in women [Chen et al., 2012; Gur et al., 1999; Ritchie et al., 2018]. Therefore, the variations investigated in the present study consistently showed associations with structural differences in specific brain regions across cortical and subcortical areas, indicating potential sex-related variations in cortical development. However, it is worth noting that the structure size may not overlap with neural activation. Notably, Kalmar *et al*. demonstrated that smaller AMY volume can facilitate increased neural activation in depressed individuals, conferring that aberrant regulation of OXT signaling can lead to psychiatric disorders [Kalmar et al., 2009].

Mounting evidence suggests that genetic variants can alter the structure and functionality of several brain regions regarding emotional and social behavior, learning, memory, motivation, language, and decision-making [Ruigrok et al., 2014]. Likewise, although we found positive and negative rsFC between different brain regions, SNPs showed analogous cognitive functions. However, interpreting positive and negative rsFC is challenging due to the functional meaning of, for instance, “weak positive rsFC” is difficult to pin down. Therefore, we used the term higher or lower rsFC instead of a positive or negative correlation, respectively. The homozygous and targeted allele carrier women had higher rsFC in temporal regions, engaging with emotional and social behavior, object and face recognition, language comprehension, and visual perception [Lewin and Herlitz, 2002; Montagne et al., 2005; Sommer et al., 2004]. While these temporal regions showed higher rsFC with a temporal region performing analogous functions in women, some areas showed additional higher rsFC with regions specialized for memory processing and integrating sensorimotor information.

Conversely, the homozygous and targeted allele carrier men indicated less rsFC than women in the occipital, parietal, and frontal regions, which process complex visual stimuli, emotion and memory regulation, and phonological processing [Pletzer et al., 2010; Proverbio et al., 2006]. Our findings showed that men had lower rsFC in processing visual (e.g., object and face recognition) and auditory information than women [Filippi et al., 2013]. However, the functional alterations for a particular cognitive function might be lower or higher depending on the presence of different allele combinations in SNPs. Nevertheless, these connectivity pairs might indicate how two brain regions communicate with each other in response to a particular event. For example, rs1042778^(–)^ men showed higher rsFC in the prefrontal cortex between IFG.oper and MFG, involving higher phonological processing, working memory, and attention allocation. Because the functional significance of this particular connectivity pair remains unknown, it proposes that men may perform better during a working memory task (e.g., Stroop task), which requires higher linguistic processing, working memory, and attention allocation. Since our study was designed to investigate rsFC differences, further investigation based on task-based functional connectivity is needed to corroborate the relationship of functional connectivity between two specific regions.

Although SC constrains functional connectivity, it has been argued that functional connectivity can produce information independently of SC [Honey et al., 2009]. Unlike the rsFC, we found that SC showed marked engagement in emotion, reward-related cognition, and behavior [Ingalhalikar et al., 2014; Joel et al., 2015]. We also found that women had higher SC primarily located in temporal and subcortical regions concerning behavior, socio-emotional and motivational salience, object and face recognition, language comprehension, visual perception, and memory processing. While rs53576^(+)^ and rs53576^(–)^ indicated some SC that denotes analogous functions, the broad dissipation at SC in women was marked in temporal and subcortical regions for the rs2254298^(+)^ and rs1042778^(–)^. This highlights that women had greater SC in several brain regions for regulating auditory and language processing (e.g., HG and STG or MTG), speech articulation, risk-reward behavior, perception, and learning memory (e.g., PU, PreCU, HI, and IN), and movement (e.g., PA). Thus, our results indicate that the rs2254298^(+)^ and rs1042778^(–)^ women are more engaged with the reward system since PU, CAU, IN, PA, HI, and TH are the components of the brain reward system. However, we acknowledge that interpreting the functional implications of SC differences can be challenging. This is because SC provides information concerning the anatomical connections between brain regions but does not directly indicate the functional activity or engagement of those regions. In addition, brain connectivity is influenced by a combination of genetic, hormonal, developmental, and environmental factors, and individual variability within each sex is substantial. Thus, further research is needed to establish the precise relationship between SC, genetic variations, and specific cognitive or behavioral functions.

Nonetheless, this structural allocation in the reward system might be augmented by fluctuations in endogenous OXT concentration. The intranasal OXT induction increased brain activation in women and decreased evoked responses in men to emotional faces [Bethlehem et al., 2013]. Because women are more engaged with endogenous OXT, they might mature a marked reward system allocation throughout brain development. Besides, other women’s hormones stimulate OXT release, up-regulate OXTR expression, and increase OXTR binding (e.g., estrogen). Thus, differences in endogenous OXT might lead to the incongruity between functional and structural connectome findings since the functionality is more dynamic than that of structural, disintegrating neural activity.

To elucidate the efficacy of endogenous OXT, we evaluated the association between endogenous OXT concentrations and functional/structural connectivity engagement with two aspects. First, we demonstrated significant sex-specific differences in the mean correlations between endogenous OXT concentrations and SC, while no significant sex-specific differences were found for rsFC. Interestingly, the rs1042778^(–)^ showed a significant association neither with SC nor rsFC. This indicates the intricate interplay between genetics, sex, and SC. Second, we performed a correlation analysis at the node level to delineate the relationship with endogenous oxytocin concentrations. Overall, the endogenous OXT correlated more with rsFC nodes than SC. Our results revealed that the variant in rs53576 was more associated with nodes of SC both in men and women. However, there were more associations with SC nodes in males with respect to variants in rs1042778. This marked association with endogenous OXT is likely to add to the broad impact of oxytocinergic systems on social behaviors, motivation, reward, desire, anxiety, and the processing of emotions.

Our findings were intrinsically limited by associations between SNPs in delineating structural and connectome-level differences. Another limitation of the present study might be the interpretation of weighted networks due to the different diffusion characteristics in white matter, especially in long-range tracks [Tsai, 2018]. However, advanced imaging techniques (e.g., diffusion microstructure) are required to uncover the factors limiting tractography [Maier-Hein et al., 2017]. In addition, we did not aim to investigate task-based functional connectivity and goal attainment, which may reveal hemispheric dominancy and underlying substrates. Epigenetic influences on the OXTR function that influence social interaction must also be considered. Therefore, there may be differential interactions with other neuromodulatory systems, such as vasopressin, opioids, steroids, and various catecholamines [Chen et al., 2020]. Further investigation is needed to characterize the mechanism of different polymorphism combinations on structure and connectivity. Finally, investigating endogenous OXT concentrations with varying polymorphism combinations might noticeably identify functional correlates of alterations in regional brain structure.

In conclusion, our data highlight the importance of considering multiple aspects of the brain morphology and structural/functional connectivity to gain a comprehensive understanding of how genetic variations and sex interact to shape the human brain, extending a large body of work on oxytocin regulation in socio-emotional processing. We show that the rs2254298^(+)^ and rs1042778^(–)^ markedly impacted structural and connectome-level functioning—especially connectome-level substrates of OXTR SNPs, which underpin socio-emotional behaviors. Moreover, by exploring the association with endogenous OXT concentrations, we provide evidence that endogenous OXT concentrations are more related to rsFC on the node level. Overall, our results contribute to our understanding of the complex relationship between genetics, brain morphology, and connectome-level substrates of OXTR SNPs.

## Data availability

The datasets collected and analyzed for the present study are available from the corresponding author upon reasonable request.

## Ethics approval

The Institutional Ethical Committee reviewed and approved the study protocol (#GO 17/877–21).

## Author contributions

KK conducted the study, developed the pipeline for data processing and analysis, analyzed and interpreted the imaging data, performed statistics, and wrote the paper; DÖ and YK conducted the study and performed ELISA and genotyping analysis, respectively. MTB supervised the genotyping analysis. EK helped with the statistical interpretation. BP and KKO contributed substantially to interpreting the data, supervised the study, and revised the manuscript critically. All authors contributed toward designing the study, helping with the grant preparation, and revising the paper, and agreed to be accountable for all aspects of the work.

## Acknowledgment

The authors thank Dr. Ethem Gelir for helping with grant preparation and Dr. Dicle Dövencioğlu for helping with MRI acquisition parameters and pilot data acquisition.

## References

Apter-Levy Y, Feldman M, Vakart A, Ebstein RP, Feldman R (2013): Impact of Maternal Depression Across the First 6 Years of Life on the Child’s Mental Health, Social Engagement, and Empathy: The Moderating Role of Oxytocin. Am J Psychiat 170:1161–1168.

Aspé-Sánchez M, Moreno M, Rivera MI, Rossi A, Ewer J (2016): Oxytocin and Vasopressin Receptor Gene Polymorphisms: Role in Social and Psychiatric Traits. Front Neurosci-switz 9:510.

Bakermans-Kranenburg MJ, IJzendoorn MH van (2008): Oxytocin receptor (OXTR) and serotonin transporter (5-HTT) genes associated with observed parenting. Soc Cogn Affect Neur 3:128–134.

Bakos J, Srancikova A, Havranek T, Bacova Z (2018): Molecular Mechanisms of Oxytocin Signaling at the Synaptic Connection. Neural Plast 2018:4864107.

Baumgartner T, Heinrichs M, Vonlanthen A, Fischbacher U, Fehr E (2008): Oxytocin Shapes the Neural Circuitry of Trust and Trust Adaptation in Humans. Neuron 58:639–650.

Behrens TEJ, Johansen-Berg H, Woolrich MW, Smith SM, Wheeler-Kingshott CAM, Boulby PA, Barker GJ, Sillery EL, Sheehan K, Ciccarelli O, Thompson AJ, Brady JM, Matthews PM (2003): Non-invasive mapping of connections between human thalamus and cortex using diffusion imaging. Nat Neurosci 6:750–757.

Behzadi Y, Restom K, Liau J, Liu TT (2007): A component based noise correction method (CompCor) for BOLD and perfusion based fMRI. Neuroimage 37:90–101.

Besson P, Dinkelacker V, Valabregue R, Thivard L, Leclerc X, Baulac M, Sammler D, Colliot O, Lehéricy S, Samson S, Dupont S (2014): Structural connectivity differences in left and right temporal lobe epilepsy. Neuroimage 100:135–144.

Bethlehem RAI, Honk J van, Auyeung B, Baron-Cohen S (2013): Oxytocin, brain physiology, and functional connectivity: A review of intranasal oxytocin fMRI studies. Psychoneuroendocrino 38:962–974.

Cahill L, Haier RJ, White NS, Fallon J, Kilpatrick L, Lawrence C, Potkin SG, Alkire MT (2001): Sex-Related Difference in Amygdala Activity during Emotionally Influenced Memory Storage. Neurobiol Learn Mem 75:1–9.

Callaert DV, Ribbens A, Maes F, Swinnen SP, Wenderoth N (2014): Assessing age-related gray matter decline with voxel-based morphometry depends significantly on segmentation and normalization procedures. Front Aging Neurosci 6:124.

Canli T, Desmond JE, Zhao Z, Gabrieli JDE (2002): Sex differences in the neural basis of emotional memories. Proc National Acad Sci 99:10789–10794.

Carson DS, Berquist SW, Trujillo TH, Garner JP, Hannah SL, Hyde SA, Sumiyoshi RD, Jackson LP, Moss JK, Strehlow MC, Cheshier SH, Partap S, Hardan AY, Parker KJ (2015): Cerebrospinal fluid and plasma oxytocin concentrations are positively correlated and negatively predict anxiety in children. Mol Psychiatr 20:1085–1090.

Carson DS, Guastella AJ, Taylor ER, McGregor IS (2013): A brief history of oxytocin and its role in modulating psychostimulant effects. J Psychopharmacol 27:231–247.

Chagnon YC, Potvin O, Hudon C, Préville M (2015): DNA methylation and single nucleotide variants in the brain-derived neurotrophic factor (BDNF) and oxytocin receptor (OXTR) genes are associated with anxiety/depression in older women. Frontiers Genetics 6:230.

Chen C-H, Gutierrez ED, Thompson W, Panizzon MS, Jernigan TL, Eyler LT, Fennema-Notestine C, Jak AJ, Neale MC, Franz CE, Lyons MJ, Grant MD, Fischl B, Seidman LJ, Tsuang MT, Kremen WS, Dale AM (2012): Hierarchical Genetic Organization of Human Cortical Surface Area. Science 335:1634–1636.

Chen X, Nishitani S, Haroon E, Smith AK, Rilling JK (2020): OXTR methylation modulates exogenous oxytocin effects on human brain activity during social interaction. Genes Brain Behav 19:e12555.

Chini B, Leonzino M, Braida D, Sala M (2014): Learning About Oxytocin: Pharmacologic and Behavioral Issues. Biol Psychiatry 76:360–366.

Dadds MR, Moul C, Cauchi A, Dobson-Stone C, Hawes DJ, Brennan J, Urwin R, Ebstein RE (2014): Polymorphisms in the oxytocin receptor gene are associated with the development of psychopathy. Dev Psychopathol 26:21–31.

Desikan RS, Ségonne F, Fischl B, Quinn BT, Dickerson BC, Blacker D, Buckner RL, Dale AM, Maguire RP, Hyman BT, Albert MS, Killiany RJ (2006): An automated labeling system for subdividing the human cerebral cortex on MRI scans into gyral based regions of interest. Neuroimage 31:968–980.

Deuse L, Wudarczyk O, Rademacher L, Kaleta P, Karges W, Kacheva S, Gründer G, Lammertz S (2018): Peripheral Oxytocin Predicts Higher-Level Social Cognition in Men Regardless of Empathy Quotient. Pharmacopsychiatry 52:148–154.

Dölen G, Darvishzadeh A, Huang KW, Malenka RC (2013): Social reward requires coordinated activity of nucleus accumbens oxytocin and serotonin. Nature 501:179–184.

Feldman R, Monakhov M, Pratt M, Ebstein RP (2016): Oxytocin Pathway Genes: Evolutionary Ancient System Impacting on Human Affiliation, Sociality, and Psychopathology. Biol Psychiat 79:174–184.

Feldman R, Zagoory-Sharon O, Weisman O, Schneiderman I, Gordon I, Maoz R, Shalev I, Ebstein RP (2012): Sensitive Parenting Is Associated with Plasma Oxytocin and Polymorphisms in the OXTR and CD38 Genes. Biol Psychiat 72:175–181.

Filippi M, Valsasina P, Misci P, Falini A, Comi G, Rocca MA (2013): The organization of intrinsic brain activity differs between genders: A resting-state fMRI study in a large cohort of young healthy subjects. Hum Brain Mapp 34:1330–1343.

Fischl B, Kouwe A van der, Destrieux C, Halgren E, Segonne F, Salat DH, Busa E, Seidman LJ, Goldstein J, Kennedy D, Caviness V, Makris N, Rosen B, Dale AM (2004): Automatically parcellating the human cerebral cortex. Cereb Cortex 14:11–22. https://www.ncbi.nlm.nih.gov/pubmed/14654453.

Fischl B, Dale AM (2000): Measuring the thickness of the human cerebral cortex from magnetic resonance images. Proc National Acad Sci 97:11050–11055.

Fischl B, Salat DH, Busa E, Albert M, Dieterich M, Haselgrove C, Kouwe A van der, Killiany R, Kennedy D, Klaveness S, Montillo A, Makris N, Rosen B, Dale AM (2002): Whole Brain Segmentation Automated Labeling of Neuroanatomical Structures in the Human Brain. Neuron 33:341–355.

Fischl B, Sereno MI, Tootell RBH, Dale AM (1999): High-resolution intersubject averaging and a coordinate system for the cortical surface. Hum Brain Mapp 8:272–284.

Furman DJ, Chen MC, Gotlib IH (2011): Variant in oxytocin receptor gene is associated with amygdala volume. Psychoneuroendocrino 36:891–897.

Gerrits NJHM, Loenhoud AC van, Berg SF van den, Berendse HW, Foncke EMJ, Klein M, Stoffers D, Werf YD van der, Heuvel OA van den (2016): Cortical Thickness, Surface Area and Subcortical Volume Differentially Contribute to Cognitive Heterogeneity in Parkinson’s Disease. Plos One 11:e0148852.

Geschwind DH, Rakic P (2013): Cortical Evolution: Judge the Brain by Its Cover. Neuron 80:633– 647.

Greve DN, Fischl B (2018): False positive rates in surface-based anatomical analysis. Neuroimage 171:6–14.

Grinevich V, Stoop R (2018): Interplay between Oxytocin and Sensory Systems in the Orchestration of Socio-Emotional Behaviors. Neuron 99:887–904.

Gur RC, Turetsky BI, Matsui M, Yan M, Bilker W, Hughett P, Gur RE (1999): Sex Differences in Brain Gray and White Matter in Healthy Young Adults: Correlations with Cognitive Performance. J Neurosci 19:4065–4072.

Hamann S, Herman RA, Nolan CL, Wallen K (2004): Men and women differ in amygdala response to visual sexual stimuli. Nat Neurosci 7:411–416.

Honey CJ, Sporns O, Cammoun L, Gigandet X, Thiran JP, Meuli R, Hagmann P (2009): Predicting human resting-state functional connectivity from structural connectivity. Proc Natl Acad Sci 106:2035–2040.

Ingalhalikar M, Smith A, Parker D, Satterthwaite TD, Elliott MA, Ruparel K, Hakonarson H, Gur RE, Gur RC, Verma R (2014): Sex differences in the structural connectome of the human brain. Proc Natl Acad Sci 111:823–828.

Inoue H, Yamasue H, Tochigi M, Abe O, Liu X, Kawamura Y, Takei K, Suga M, Yamada H, Rogers MA, Aoki S, Sasaki T, Kasai K (2010): Association Between the Oxytocin Receptor Gene and Amygdalar Volume in Healthy Adults. Biol Psychiat 68:1066–1072.

Israel S, Lerer E, Shalev I, Uzefovsky F, Reibold M, Bachner-Melman R, Granot R, Bornstein G, Knafo A, Yirmiya N, Ebstein RP (2008): Molecular genetic studies of the arginine vasopressin 1a receptor (AVPR1a) and the oxytocin receptor (OXTR) in human behaviour: from autism to altruism with some notes in between. Prog Brain Res 170:435–449.

Jacob S, Brune CW, Carter CS, Leventhal BL, Lord C, Cook EH (2007): Association of the oxytocin receptor gene (OXTR) in Caucasian children and adolescents with autism. Neurosci Lett 417:6–9.

Jenkinson M, Beckmann CF, Behrens TEJ, Woolrich MW, Smith SM (2012): FSL. Neuroimage 62:782–790.

Joel D, Berman Z, Tavor I, Wexler N, Gaber O, Stein Y, Shefi N, Pool J, Urchs S, Margulies DS, Liem F, Hänggi J, Jäncke L, Assaf Y (2015): Sex beyond the genitalia: The human brain mosaic. Proc Natl Acad Sci 112:15468–15473.

Jurek B, Neumann ID (2018): The Oxytocin Receptor: From Intracellular Signaling to Behavior. Physiol Rev 98:1805–1908.

Kalmar JH, Wang F, Chepenik LG, Womer FY, Jones MM, Pittman B, Shah MP, Martin A, Constable RT, Blumberg HP (2009): Relation Between Amygdala Structure and Function in Adolescents With Bipolar Disorder. J Am Acad Child Adolesc Psychiatry 48:636–642.

Kennedy KM, Erickson KI, Rodrigue KM, Voss MW, Colcombe SJ, Kramer AF, Acker JD, Raz N (2009): Age-related differences in regional brain volumes: A comparison of optimized voxel-based morphometry to manual volumetry. Neurobiol Aging 30:1657–1676.

Kirsch P, Esslinger C, Chen Q, Mier D, Lis S, Siddhanti S, Gruppe H, Mattay VS, Gallhofer B, Meyer-Lindenberg A (2005): Oxytocin Modulates Neural Circuitry for Social Cognition and Fear in Humans. J Neurosci 25:11489–11493.

Kosfeld M, Heinrichs M, Zak PJ, Fischbacher U, Fehr E (2005): Oxytocin increases trust in humans. Nature 435:673–676.

Lewin C, Herlitz A (2002): Sex differences in face recognition—Women’s faces make the difference. Brain Cogn 50:121–128.

Ludwig M, Leng G (2006): Dendritic peptide release and peptide-dependent behaviours. Nat Rev Neurosci 7:126–136.

Luo S, Zhu Y, Fan L, Gao D, Han S (2020): Resting-state brain network properties mediate the association between the oxytocin receptor gene and interdependence. Soc Neurosci 15:296– 310.

Maier-Hein KH, Neher PF, Houde J-C, Côté M-A, Garyfallidis E, Zhong J, Chamberland M, Yeh F-C, Lin Y-C, Ji Q, Reddick WE, Glass JO, Chen DQ, Feng Y, Gao C, Wu Y, Ma J, He R, Li Q, Westin C-F, Deslauriers-Gauthier S, González JOO, Paquette M, St-Jean S, Girard G, Rheault F, Sidhu J, Tax CMW, Guo F, Mesri HY, Dávid S, Froeling M, Heemskerk AM, Leemans A, Boré A, Pinsard B, Bedetti C, Desrosiers M, Brambati S, Doyon J, Sarica A, Vasta R, Cerasa A, Quattrone A, Yeatman J, Khan AR, Hodges W, Alexander S, Romascano D, Barakovic M, Auría A, Esteban O, Lemkaddem A, Thiran J-P, Cetingul HE, Odry BL, Mailhe B, Nadar MS, Pizzagalli F, Prasad G, Villalon-Reina JE, Galvis J, Thompson PM, Requejo FDS, Laguna PL, Lacerda LM, Barrett R, Dell’Acqua F, Catani M, Petit L, Caruyer E, Daducci A, Dyrby TB, Holland-Letz T, Hilgetag CC, Stieltjes B, Descoteaux M (2017): The challenge of mapping the human connectome based on diffusion tractography. Nat Commun 8:1349.

Maud C, Ryan J, McIntosh JE, Olsson CA (2018): The role of oxytocin receptor gene (OXTR) DNA methylation (DNAm) in human social and emotional functioning: a systematic narrative review. Bmc Psychiatry 18:154.

McQuaid RJ, McInnis OA, Abizaid A, Anisman H (2014): Making room for oxytocin in understanding depression. Neurosci Biobehav Rev 45:305–322.

Meyer-Lindenberg A (2008): Impact of prosocial neuropeptides on human brain function. Prog Brain Res 170:463–470.

Meyer-Lindenberg A, Domes G, Kirsch P, Heinrichs M (2011): Oxytocin and vasopressin in the human brain: social neuropeptides for translational medicine. Nat Rev Neurosci 12:524–538.

Mitre M, Marlin BJ, Schiavo JK, Morina E, Norden SE, Hackett TA, Aoki CJ, Chao MV, Froemke RC (2016): A Distributed Network for Social Cognition Enriched for Oxytocin Receptors. J Neurosci 36:2517–2535.

Montagne B, Kessels RPC, Frigerio E, Haan EHF de, Perrett DI (2005): Sex differences in the perception of affective facial expressions: Do men really lack emotional sensitivity? Cogn Process 6:136–141.

Onitsuka T, Shenton ME, Salisbury DF, Dickey CC, Kasai K, Toner SK, Frumin M, Kikinis R, Jolesz FA, McCarley RW (2004): Middle and Inferior Temporal Gyrus Gray Matter Volume Abnormalities in Chronic Schizophrenia: An MRI Study. Am J Psychiat 161:1603–1611.

Panizzon MS, Fennema-Notestine C, Eyler LT, Jernigan TL, Prom-Wormley E, Neale M, Jacobson K, Lyons MJ, Grant MD, Franz CE, Xian H, Tsuang M, Fischl B, Seidman L, Dale A, Kremen WS (2009): Distinct Genetic Influences on Cortical Surface Area and Cortical Thickness. Cereb Cortex 19:2728–2735.

Pekarek BT, Hunt PJ, Arenkiel BR (2020): Oxytocin and Sensory Network Plasticity. Front Neurosci-switz 14:30.

Plasencia G, Luedicke JM, Nazarloo HP, Carter CS, Ebner NC (2019): Plasma oxytocin and vasopressin levels in young and older men and women: Functional relationships with attachment and cognition. Psychoneuroendocrino 110:104419.

Pletzer B, Kerschbaum H, Klimesch W (2010): When frequencies never synchronize: The golden mean and the resting EEG. Brain Res 1335:91–102.

Proverbio AM, Brignone V, Matarazzo S, Zotto MD, Zani A (2006): Gender differences in hemispheric asymmetry for face processing. BMC Neurosci 7:44.

Quintana DS, Rokicki J, Meer D van der, Alnæs D, Kaufmann T, Córdova-Palomera A, Dieset I, Andreassen OA, Westlye LT (2019): Oxytocin pathway gene networks in the human brain. Nat Commun 10:668.

Rajmohan V, Mohandas E (2007): The limbic system. Indian J Psychiatry 49:132–139.

Ritchie SJ, Cox SR, Shen X, Lombardo MV, Reus LM, Alloza C, Harris MA, Alderson HL, Hunter S, Neilson E, Liewald DCM, Auyeung B, Whalley HC, Lawrie SM, Gale CR, Bastin ME, McIntosh AM, Deary IJ (2018): Sex Differences in the Adult Human Brain: Evidence from 5216 UK Biobank Participants. Cereb Cortex 28:2959–2975.

Rodrigues SM, Saslow LR, Garcia N, John OP, Keltner D (2009): Oxytocin receptor genetic variation relates to empathy and stress reactivity in humans. Proc National Acad Sci 106:21437–21441.

Ruigrok ANV, Salimi-Khorshidi G, Lai M-C, Baron-Cohen S, Lombardo MV, Tait RJ, Suckling J (2014): A meta-analysis of sex differences in human brain structure. Neurosci Biobehav Rev 39:34–50.

Savaskan E, Ehrhardt R, Schulz A, Walter M, Schächinger H (2008): Post-learning intranasal oxytocin modulates human memory for facial identity. Psychoneuroendocrino 33:368–374.

Sicorello M, Dieckmann L, Moser D, Lux V, Luhmann M, Schlotz W, Kumsta R (2020): Oxytocin and the stress buffering effect of social company: A genetic study in daily life. Soc Cogn Affect Neur 15:293–301.

Sommer IEC, Aleman A, Bouma A, Kahn RS (2004): Do women really have more bilateral language representation than men? A meta-analysis of functional imaging studies. Brain 127:1845–1852.

Strauss GP, Chapman HC, Keller WR, Koenig JI, Gold JM, Carpenter WT, Buchanan RW (2019): Endogenous oxytocin levels are associated with impaired social cognition and neurocognition in schizophrenia. J Psychiatr Res 112:38–43.

Tops M, Huffmeijer R, Linting M, Grewen KM, Light KC, Koole SL, Bakermans-Kranenburg MJ, IJzendoorn MH van (2013): The role of oxytocin in familiarization-habituation responses to social novelty. Front Psychol 4:761.

Tost H, Kolachana B, Hakimi S, Lemaitre H, Verchinski BA, Mattay VS, Weinberger DR, Meyer–Lindenberg A (2010): A common allele in the oxytocin receptor gene (OXTR) impacts prosocial temperament and human hypothalamic-limbic structure and function. Proc National Acad Sci 107:13936–13941.

Tournier JD, Calamante F, Connelly A (2010): Improved probabilistic streamlines tractography by 2nd order integration over fibre orientation distributions. In: Proceedings of the international society for magnetic resonance in medicine. Ismrm. Vol. 1670, p.

Tournier J-D, Calamante F, Connelly A (2007): Robust determination of the fibre orientation distribution in diffusion MRI: Non-negativity constrained super-resolved spherical deconvolution. Neuroimage 35:1459–1472.

Tournier J -Donald, Calamante F, Connelly A (2013): Determination of the appropriate b value and number of gradient directions for high-angular-resolution diffusion-weighted imaging. Nmr Biomed 26:1775–1786.

Tournier J-D, Smith R, Raffelt D, Tabbara R, Dhollander T, Pietsch M, Christiaens D, Jeurissen B, Yeh C-H, Connelly A (2019): MRtrix3: A fast, flexible and open software framework for medical image processing and visualisation. Neuroimage 202:116137.

Tsai S-Y (2018): Reproducibility of structural brain connectivity and network metrics using probabilistic diffusion tractography. Sci Rep-uk 8:11562.

Waller R, Corral-Frías NS, Vannucci B, Bogdan R, Knodt AR, Hariri AR, Hyde LW (2016): An oxytocin receptor polymorphism predicts amygdala reactivity and antisocial behavior in men. Soc Cogn Affect Neur 11:1218–1226.

Weisman O, Zagoory-Sharon O, Schneiderman I, Gordon I, Feldman R (2013): Plasma oxytocin distributions in a large cohort of women and men and their gender-specific associations with anxiety. Psychoneuroendocrino 38:694–701.

Whitfield-Gabrieli S, Nieto-Castanon A (2012): Conn: A Functional Connectivity Toolbox for Correlated and Anticorrelated Brain Networks. Brain Connectivity 2:125–141.

Winkler AM, Kochunov P, Blangero J, Almasy L, Zilles K, Fox PT, Duggirala R, Glahn DC (2010): Cortical thickness or grey matter volume? The importance of selecting the phenotype for imaging genetics studies. Neuroimage 53:1135–1146.

Yamasue H, Yee JR, Hurlemann R, Rilling JK, Chen FS, Meyer-Lindenberg A, Tost H (2012): Integrative Approaches Utilizing Oxytocin to Enhance Prosocial Behavior: From Animal and Human Social Behavior to Autistic Social Dysfunction. J Neurosci 32:14109–14117a.

Yoshida M, Takayanagi Y, Inoue K, Kimura T, Young LJ, Onaka T, Nishimori K (2009): Evidence That Oxytocin Exerts Anxiolytic Effects via Oxytocin Receptor Expressed in Serotonergic Neurons in Mice. J Neurosci 29:2259–2271.

Zalesky A, Fornito A, Bullmore ET (2010): Network-based statistic: Identifying differences in brain networks. Neuroimage 53:1197–1207.

Zhang T-Y, Shahrokh D, Hellstrom IC, Wen X, Diorio J, Breuillaud L, Caldji C, Meaney MJ (2020): Brain-Derived Neurotrophic Factor in the Nucleus Accumbens Mediates Individual Differences in Behavioral Responses to a Natural, Social Reward. Mol Neurobiol 57:290–301.

Zik JB, Roberts DL (2015): The many faces of oxytocin: Implications for psychiatry. Psychiat Res 226:31–37.

