## Supplementary Data for "Situating the oxytocin receptor gene polymorphisms in the context of structural and connectome-level substrates and association with endogenous oxytocin"

[1] Department of Physiology, Faculty of Medine, Hacettepe University, Ankara, Türkiye
[2] Department of Physiology, Faculty of Medine, Balıkesir University, Balıkesir, Türkiye
[3] Department of Physiology, Faculty of Medine, Kırklareli University, Kırklareli, Türkiye
[4] Department of Biostatistics, Faculty of Medine, Hacettepe University, Ankara, Türkiye
[6] Department of Radiology, Division of Neuroradiology, University of California Davis Medical Center, Sacramento, CA, USA

**Table S1.** Significant sex-specific cortical surface area (cSA) differences

| cSA ($mm^2$) | | mean ± SD | | | |
|---|---|---|---|---|---|
| **SNP** | **Region** | **Women** | **Men** | **Cluster size** | **CWP** |
| rs53576$^{(+)}$ | L MTG | 0.73 ± 0.10 | 0.74 ± 0.13 | 405.62 | 0.04 |
| | L FG | 0.76 ± 0.11 | 0.80 ± 0.10 | 403.81 | 0.04 |
| rs1042778$^{(+)}$ | R ITG | 0.89 ± 0.14 | 0.99 ± 0.20 | 489.73 | 0.03 |
| rs2254298$^{(+)}$ | L MTG | 0.79 ± 0.10 | 0.87 ± 0.13 | 2336.68 | < 0.001 |
| | L PreCG | 0.47 ± 0.06 | 0.49 ± 0.09 | 1345.92 | < 0.001 |
| | L SPG | 0.51 ± 0.09 | 0.53 ± 0.14 | 760.14 | 0.01 |
| | R ITG | 0.80 ± 0.11 | 0.86 ± 0.14 | 1824.75 | < 0.001 |
| | R SFG | 0.87 ± 0.09 | 0.91 ± 0.12 | 662.1 | 0.02 |
| | R PreCG | 0.46 ± 0.06 | 0.47 ± 0.07 | 463.39 | 0.04 |
| rs53576$^{(–)}$ | L PreCG | 0.47 ± 0.07 | 0.52 ± 0.10 | 2300.75 | < 0.001 |
| | L LOC | 0.85 ± 0.11 | 0.83 ± 0.11 | 622.70 | 0.01 |
| | L SPG | 0.50 ± 0.10 | 0.54 ± 0.10 | 616.71 | 0.01 |
| | L ITG | 0.81 ± 0.10 | 0.85 ± 0.15 | 366.06 | 0.04 |
| | R SPG | 0.77 ± 0.13 | 0.76 ± 0.09 | 844.76 | 0.01 |
| | R PreCG | 0.49 ± 0.10 | 0.50 ± 0.10 | 628.73 | 0.03 |
| rs1042778$^{(–)}$ | L LOC | 0.78 ± 0.10 | 0.82 ± 0.10 | 1618.06 | < 0.001 |
| | L PreCG | 0.51 ± 0.08 | 0.50 ± 0.09 | 629.44 | 0.02 |
| | L MTG | 0.73 ± 0.11 | 0.77 ± 0.09 | 438.46 | 0.04 |
| | R CU | 0.83 ± 0.12 | 0.84 ± 0.12 | 824.79 | 0.01 |
| rs2254298$^{(–)}$ | L FG | 0.79 ± 0.09 | 0.89 ± 0.11 | 723.33 | 0.01 |
| | R CU | 0.81 ± 0.09 | 0.89 ± 0.12 | 898.11 | 0.005 |

L, left; R, right; SNP, single nucleotide polymorphism; SNP$^{(+)}$, homozygous allele; SNP$^{(–)}$, targeted allele carriers; MTG, middle temporal gyrus; FG, fusiform gyrus; ITG, inferior temporal gyrus; PreCG, precentral gyrus; SPG, superior parietal gyrus; SFG, superior frontal gyrus; LOC, lateral occipital cortex; CWP, cluster-wise *p*-value; SD, standard deviation.

**Table S2.** Significant sex-specific cortical gray matter (GM) volume differences

| Cortical GM volume ($mm^3$) | | mean ± SD | | | |
|---|---|---|---|---|---|
| SNP | Region | Women | Men | Cluster size | CWP |
| rs53576$^{(+)}$ | L ITG | 3.09 ± 0.49 | 3.32 ± 0.69 | 352.83 | 0.01 |
| rs2254298$^{(+)}$ | L ITG | 2.29 ± 0.42 | 2.53 ± 0.57 | 438.31 | 0.007 |
| | R SFG | 2.69 ± 0.30 | 2.88 ± 0.50 | 307.81 | 0.02 |
| | R LOC | 1.88 ± 0.40 | 2.06 ± 0.46 | 293.17 | 0.02 |
| | R FG | 2.31 ± 0.28 | 2.43 ± 0.48 | 227.75 | 0.04 |
| rs53576$^{(-)}$ | L PreCG | 1.06 ± 0.20 | 1.22 ± 0.24 | 687.31 | 0.03 |
| | L ITG | 2.00 ± 0.32 | 2.13 ± 0.45 | 246.46 | 0.02 |
| | L SFG | 2.03 ± 0.50 | 2.30 ± 0.56 | 202.85 | 0.04 |
| | R SPG | 1.68 ± 0.33 | 1.70 ± 0.25 | 647.57 | 0.001 |
| | R PreCG | 1.38 ± 0.36 | 1.44 ± 0.33 | 261.25 | 0.03 |
| rs1042778$^{(-)}$ | L SFG | 2.21 ± 0.55 | 2.13 ± 0.57 | 246.57 | 0.03 |
| | R SPG | 1.86 ± 0.42 | 1.76 ± 0.37 | 379.36 | 0.007 |
| rs2254298$^{(-)}$ | L FG | 2.12 ± 0.32 | 2.49 ± 0.37 | 403.06 | 0.002 |
| | L PreCU | 1.24 ± 0.29 | 1.43 ± 0.29 | 192.03 | 0.03 |
| | R PreCU | 1.51 ± 0.24 | 1.72 ± 0.30 | 191.67 | 0.03 |

L, left; R, right; SNP, single nucleotide polymorphism; SNP$^{(+)}$, homozygous allele; SNP$^{(-)}$, targeted allele carriers; ITG, inferior temporal gyrus; SFG, superior frontal gyrus; LOC, lateral occipital cortex; PreCG, precentral gyrus; SPG, superior parietal gyrus; FG, fusiform gyrus; PreCU, precuneus; CWP, cluster-wise *p*-value; SD, standard deviation.

**Table S3.** Significant sex-specific subcortical gray matter (GM) volume differences

| Subcortical (GM) volume ($mm^3$) | | median (IQR) | | | |
|---|---|---|---|---|---|
| **SNP** | **Region** | **Women** | **Men** | ***U*** | $p_{corrected}$ |
| rs53576$^{(+)}$ | L AMY | 1347 (183) | 1500 (137) | 43 | 0.006 |
| rs1042778$^{(+)}$ | R AMY | 1421 (249) | 1664 (387) | 14 | 0.001 |
| rs2254298$^{(+)}$ | L AMY | 1369 (294) | 1521 (325) | 99 | 0.003 |
| | L CAU | 584 (153) | 676 (187) | 102 | 0.004 |
| | L vDC | 3735 (41) | 4159 (515) | 78 | < 0.001 |
| rs53576$^{(–)}$ | L AMY | 1329 (277) | 1606 (431) | 51 | 0.005 |
| | R PU | 5031 (520) | 5629 (623) | 49 | 0.003 |
| rs1042778$^{(–)}$ | L AMY | 1310 (227) | 1485 (223) | 71 | 0.001 |
| | L HI | 3818 (338) | 4287 (553) | 67 | 0.001 |

L, left; R, right; SNP, single nucleotide polymorphism; SNP$^{(+)}$, homozygous allele; SNP$^{(–)}$, targeted allele carriers; AMY, amygdala; CAU, caudate; vDC, ventral diencephalon; PU, putamen; HI, hippocampus; IQR, interquartile range. The *p*-value was Bonferroni corrected ($p_{corrected}$).

**Table S4.** Significant structural connectivity (SC) pairs after performing network-based statistics

| SNP | node 1 | node 2 | $t$ | $p_{corrected}$ |
|---|---|---|---|---|
| rs53576(+) | R HI | L PostCG | 3.71 | 0.01 |
| | R HI | L PreCG | 3.72 | |
| | R HG | L IN | 3.81 | |
| | R HG | L HI | 4.09 | |
| rs2254298(+) | L STG | R STG | 4.13 | 0.002 |
| | L STG | R HG | 5.66 | |
| | L HG | L TH | 3.74 | |
| | L HG | R PU | 4.33 | |
| | L HG | R PreCU | 4.11 | |
| | L HG | R STG | 4.17 | |
| | L HG | R HG | 4.52 | |
| | L HG | R IN | 3.78 | |
| | L IN | R STG | 3.97 | |
| | L IN | R HG | 3.98 | |
| | L HI | R PU | 4.19 | |
| | L HI | R PA | 3.73 | |
| | L HI | R HG | 4.24 | |
| | R HG | L MTG | 3.74 | |
| | R HG | L PU | 3.77 | |
| rs53576(–) | L EC | R PA | 4.35 | 0.04 |
| | L EC | R AMY | 4.90 | |
| | L PaHG | R PA | 4.71 | |

**Table S4.** *Cont.*

| SNP | node 1 | node 2 | $t$ | $p_{corrected}$ |
|---|---|---|---|---|
| rs1042778$^{(-)}$ | L HI | R CAU | 3.99 | 0.002 |
| | L HI | R PU | 3.78 | |
| | L HI | R PA | 3.72 | |
| | L HI | R HI | 3.72 | |
| | L HI | R HG | 3.73 | |
| | R HI | L CAU | 4.05 | |
| | R HI | L PU | 3.83 | |
| | R HI | L PA | 3.82 | |
| | R HG | L STG | 3.95 | |
| | R HG | L IN | 3.84 | |
| | R HG | R PreCU | 3.74 | |
| | R PreCG | L PaHG | 3.76 | |

L, left; R, right; SNP, single nucleotide polymorphism; SNP$^{(+)}$, homozygous allele; SNP$^{(-)}$, targeted allele carriers; HI, hippocampus; HG, Heschl's gyrus; PostCG, postcentral gyrus; PreCG, precentral gyrus; IN, insula; STG, superior temporal gyrus; TH, thalamus; PU, putamen; PreCU, precuneus; PA, pallidum; MTG, middle temporal gyrus; CAU, caudate; PaHG, parahippocampal gyrus; EC, entorhinal cortex; AMY, amygdala. The $p$-value was a family-wise error (FWE) corrected ($p_{corrected}$).
